# CKS1-dependent proteostatic regulation has dual roles combating acute myeloid leukemia whilst protecting normal hematopoiesis

**DOI:** 10.1101/2020.12.27.423419

**Authors:** W. Grey, A. Rio-Machin, P. Casado-Izquierdo, J.J. Miettinen, F. Copley, A. Parsons, C.A. Heckman, P. Cutillas, J. Gribben, J. Fitzgibbon, D. Bonnet

**Affiliations:** Haematopoietic Stem Cell Laboratory, The Francis Crick Institute, London, U.K.; Centre for Genomics and Computational Biology, Barts Cancer Institute, London, U.K.; Cell signalling and proteomics group, Centre for Genomics and Computational Biology, Barts Cancer Institute, London, U.K.; Institute for Molecular Medicine Finland – FINN, HiLIFE – Helsinki Institute of Life Science, iCAN Digital Precision Cancer Medicine Flagship, University of Helsinki, Helsinki, Finland; Centre for Haemato-Oncology, Barts Cancer Institute, London, U.K.

**Keywords:** CKS1, Proteostasis, AML, Hematopoiesis, Chemotherapy

## Abstract

Acute myeloid leukemia (AML) is an aggressive hematological disorder comprising a hierarchy of quiescent leukemic stem cells (LSCs) and proliferating blasts with limited self-renewal ability. AML has a dismal prognosis, with extremely low two-year survival rates in the poorest cytogenetic risk patients, primarily due to the failure of intensive chemotherapy protocols unable to deplete LSCs, which reconstitute the disease *in vivo*, and the significant toxicity towards healthy hematopoietic cells. Whilst much work has been done to identify genetic and epigenetic vulnerabilities in AML LSCs, little is known about protein dynamics and the role of protein degradation in drug resistance and relapse. Here, using a highly specific inhibitor of the SCF^SKP2-CKS1^ complex, we report a dual role for CKS1-dependent protein degradation in reducing AML blasts *in vivo*, and importantly depleting LSCs. Whilst many AML LSC targeted therapies show significant toxicity to healthy hematopoiesis, inhibition of CKS1-dependent protein degradation has the opposite effect, protecting normal hematopoietic cells from chemotherapeutic toxicity. Together these findings demonstrate CKS1-dependent proteostasis is key for normal and malignant hematopoiesis.

**Significance:** CKS1-dependent protein degradation is a specific vulnerability in AML LSCs. Specific inhibition of SCF^SKP2-CKS1^ is lethal to *CKS1B*^*high*^ AML blasts and all AML LSCs. Normal hematopoiesis is protected from chemotherapeutic toxicity by inhibition of CKS1-dependent protein degradation, substantiating a dual role for CKS1-dependent protein degradation in clinical treatment of AML.

## Introduction

Acute myeloid leukaemia (AML) is a heterogeneous, aggressive disease of the hematopoietic system, arising from hematopoietic stem/progenitor cells. In recent years, several reports have demonstrated that the current approach (induction chemotherapy) and new protocols (epigenetic targeting) still have severe limitations due to significant plasticity in the AML epigenome, metabolic adaptions and the presence of drug resistance leukemic stem cells (LSCs). With the average two-year survival rate as low as 5-15% in poor risk, older patients (>65yr), there is an unmet critical need for new therapeutic approaches(1). Recent developments, targeting the anti-apoptotic protein BCL2, has demonstrated that therapies affecting protein networks holds great promise for the poorest prognosis AMLs(2,3), yet resistance still emerges for a subset of patients through mitochondrial adaptions in residual leukemic cells(4,5).

New approaches such as BCL2 targeting in combination with classical induction chemotherapy still hold excellent promise, but a critical failure is still the severe off-target toxicity produced by induction chemotherapy protocols and new targeted therapies alike(6). Indeed, reducing blast count with cytarabine/doxorubicin treatment severely affects normal hematopoietic progenitor cells, stressing the hematopoietic system(7). Whilst bone marrow transplantation remains the gold standard consolidation therapy in AML(8), boosting normal hematopoiesis to outcompete residual AML, combined with a reduction in severe cytopenia immediately after chemotherapy, would be beneficial to overall survival.

A better understanding of the biological differences between normal and malignant hematopoietic cells is needed to achieve selective AML targeting, without toxicity to normal cells. We previously reported a proteostatic axis between the cyclin-dependent kinase subunits Cks1 and Cks2, and the mixed lineage leukaemia 1 protein (Mll1), a key protein hijacked during neoplastic transformation of the hematopoietic system(9). Cks1 and Cks2 have many overlapping and independent roles in balancing protein homeostasis (proteostasis) throughout the cell cycle, ensuring correct G0/G1 transition(10), chromatin separation(11–13) and DNA repair(10,14,15). Whilst it was originally thought Cks1 and Cks2 function solely through CDK-dependent activities(16–18), CDK-independent functions were later reported, in concert with the SCF^SKP2^ and APC^CDC20^ E3 ubiquitin ligase complexes, important for selective protein degradation(10,11,19).

The many functions of Cks1 and Cks2 place this axis at the centre of normal cell growth and development, and potentially central to AML development. Here, we investigated the role of CKS1-dependent protein degradation in a poor risk AML cohort with few treatment options. We explored the therapeutic potential in *CKS1B* expressing poor risk AML and demonstrated high efficacy in reducing total AML burden in *CKS1B*^*high*^ expressing samples. Critically, in both *CKS1B*^*high* and *low*^ AML we demonstrate a significant reduction of the chemotherapy refractory LSCs. In contrast, CKS1 inhibition has the opposite effect on normal hematopoiesis, improving stem cell functionality and conferring protection from chemotherapeutic toxicity.

## Results

### *CKS1B* expression dictates the susceptibility of AML to specific chemotherapy

The expression of *CKS1B* varies in normal hematopoiesis, with moderate expression in hematopoietic stem cells (HSCs), the highest expression in myeloid progenitors, and the lowest expression in terminally differentiated cells (Supp. Fig. 1A). We previously reported *CKS1B* upregulation in *MLL1*-rearranged AML, acute lymphoblastic leukemia (ALL) and chronic myeloid leukemia (CML) compared to peripheral blood mononuclear cells (PBMCs)(9). This trend of *CKS1B* expression is conserved amongst most AML cytogenetic subtypes (Supp. Fig. 1A), despite high variability of *CKS1B* expression within AML subtypes, with large standard deviations compared to one of its key upstream proteostatic regulation partners *SKP2* (Supp. Fig. 1B). Whilst *CKS1B* expression can be prognostic in a variety of cancers(20–22), it is not prognostic in AML at the RNA level (Supp. Fig. 1C).

Considering that the key function of CKS1 is substrate recognition, and adapting the binding of phosphoproteins to their kinase or ubiquitin ligase regulators(18,19), we hypothesized that high *CKS1B* AMLs may show a selective susceptibility to inhibition of CKS1-dependent protein degradation (SCF^SKP2-CKS1^ E3 ligase inhibitor), and inhibition of associated signalling pathways. To address this key question, we screened a cohort of cytogenetically poor risk AMLs, spanning a variety of FAB and molecular subtypes, with a broad range of clinically approved and early development phase compounds and correlated drug sensitivity with *CKS1B* expression (Figure 1A-B, Supp. Table 1).

**Figure 1.**
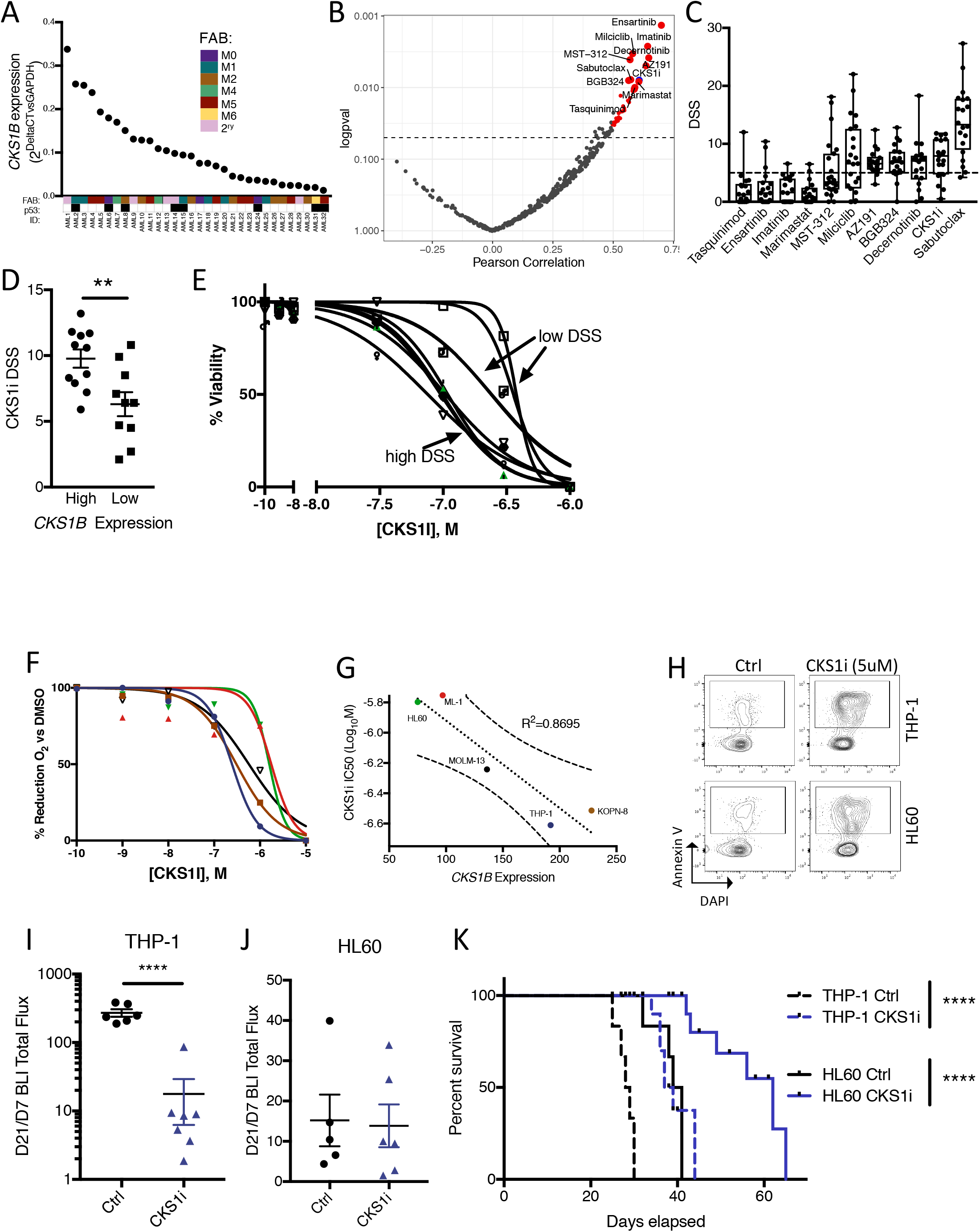
*CKS1B* expression status can predict sensitivity to inhibition of CKS1-dependent protein degradation in AML. **A.** Expression of *CKS1B* (relative to *GAPDH*) in a poor risk AML cohort. FAB and p53 status are indicated for each patient (FAB color coded, p53 status: white = WT; black = mutant; N=32). **B.** Pearson’s correlation of *CKS1B* expression versus DSS for a panel of clinically available and investigational compounds. Red indicates significant correlations above R^2^=0.5. Blue indicates CKS1i (N=21). **C.** Diagram of action for CKS1i binding and inhibition of the SCF^SKP2-CKS1^ ubiquitin ligase complex **D.** Percentage of viability of example patient AML samples cultured for 72 hours with indicated doses of CKS1i. **E.** Allocation of patient AML in high or low *CKS1B* expression (50^th^ percentile) and compared for CKS1i DSS (N=21). **F.** Percentage viability of AML cell lines cultured for 48 hours with indicated doses of CKS1i (N=3 for all cell lines on graph). **G.** Correlation between AML cell line CKS1i IC_50_ and *CKS1B* expression. 95% confidence intervals presented. Pearson’s correlation coefficient was calculated for correlation (R^2^). **H.** Representative FACS plots for induction of apoptosis in the indicated AML cell lines by presence of annexin V at the cell surface in response to CKS1i (5μM) at 48 hours. Fold change *in vivo* leukemic burden of **I.** THP-1 (Ctrl N=6, CKS1i N=7) and **J.** HL60 (Ctrl N=5, CKS1i N=6) cells day 21 (9 days post-CKS1i) versus day 7 (pre-CKS1i) expressed as bioluminescent total flux intensity. **K.** Overall survival of xenografts carrying THP-1 and HL60 cell lines control or treated with CKS1i. A Student’s *t*-test was used to calculate significance of difference for all graphs unless otherwise stated. ** p<0.005; **** p<0.0001.

*CKS1B* expression significantly correlated with response to 52 individual compounds including classical DNA damaging chemotherapy (e.g. Cisplatin, R=0.43 p=0.04), kinase inhibitors (e.g. Ensartinib, R=0.70, p=0.001), BCL2 family inhibitors (e.g. Sabutoclax, R=0.58, p=0.008) and the SCF^SKP2-CKS1^ E3 ligase inhibitor (hereafter referred to as CKS1i; R=0.61, p=0.008; Figure 1B-C, Supp. Fig. 1D). Susceptibility to CKS1i significantly correlated with *CKS1B* expression overall (Supp. Fig. 1D), and especially well in complex karyotype patients (Supp. Fig. 1E). Separating patients at the 50^th^ percentile by *CKS1B* expression revealed significantly increased drug sensitivity in *CKS1B*^*high*^ AML versus *CKS1B*^*low*^ AML patients (Figure 1D-E), indicating that RNA expression of *CKS1B* can be a selection criterion for targeting SCF^SKP2-CKS1^ dependent protein degradation in AML.

The susceptibility of patient AML samples to CKS1i based on *CKS1B* expression is well conserved in cell lines used to study molecular characteristics of AML. Dose dependent response (IC_50_) was lower in *CKS1B*^*high*^ AML cell lines THP-1 (*MLL-AF9*) and KOPN-8 (*MLL-ENL*) than in *CKS1B*^*low*^ cell lines ML-1 (*MLL-AF6*) and HL60 (*cMyc*, Figure 1F) and IC_50_ values correlate well with *CKS1B* expression (Figure 1G). At higher doses, CKS1i induces cell death in both high and low *CKS1B* cell lines (5μM; Figure 1H), a dose permissive to healthy cells *in vitro*(9). Susceptibility of AML cell lines to CKS1i is maintained *in vivo* accordingly with *CKS1B* expression. *CKS1B*^high^ THP-1 and *CKS1B*^low^ HL60 cells engrafted in immunodeficient mice (NSG) showed similarly diverging sensitivity to a single course of CKS1i treatment (10mg/kg, 5 days I.P.). THP-1 leukemic burden in NSG mice was significantly reduced by CKS1i treatment (Figure 1I), whereas in HL60, leukemic burden remained similar between control and CKS1i treated mice (Figure 1J). Interestingly, despite the difference in overall tumor burden after chemotherapy, both xenograft models showed significant overall improved survival when treated with CKS1i compared to controls (Figure 1K).

In order to investigate the effect of CKS1i on primary patient AML *in vivo*, we selected five primary patient samples with a range of *CKS1B* expression (Figure 2A, Supp. Table 1), for which we have previously reported robust engraftment in NSG mice(23,24) (Figure 2B). A single course of CKS1i (10mg/kg, 5 days I.P.) was administered and significantly reduced the AML burden in patients with the highest *CKS1B* expression (AML12 and AML21). A trend towards reduced AML burden was seen at an intermediate level of *CKS1B* expression (AML26), but had no significant effect was seen on patients with the lowest *CKS1B* expression (AML27 and AML32). As such, *CKS1B* expression levels correlated well with outcome (R=−0.446; Figure 2C, Supp. Fig 2A). In agreement with our observations of AML cell line *in vivo* responses, all CKS1i treated AML xenografts showed significantly improved overall survival compared to untreated controls (Figure 2D), indicating that overall survival conferred by CKS1i was not only due to changes in bulk leukemic burden.

**Figure 2.**
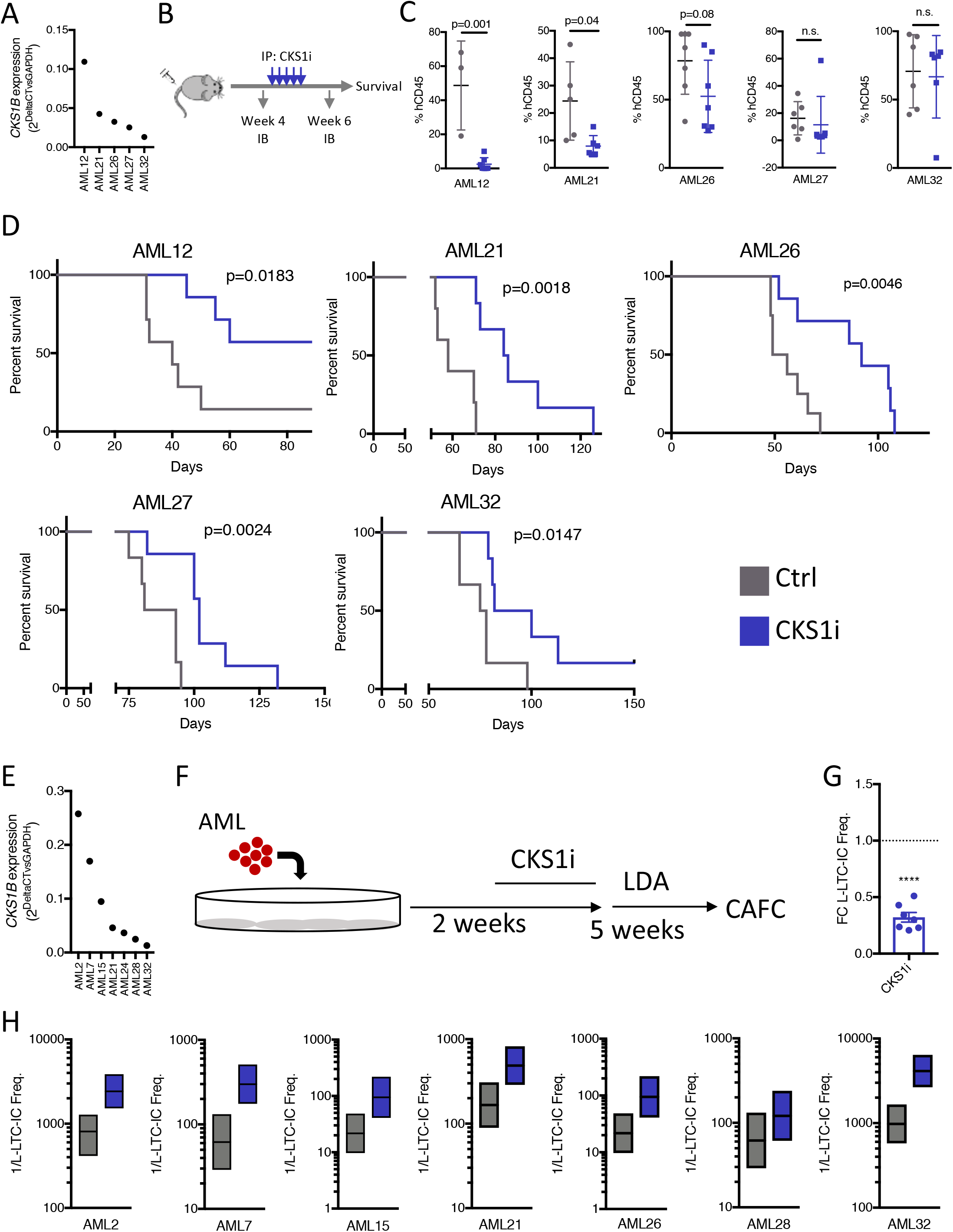
Inhibition of CKS1-dependent protein degradation selectively kills bulk AML *in vivo* and depletes leukemic stem cell potential. **A.** *CKS1B* expression (relative to *GAPDH*) for patient AMLs tested *in vivo* (N=5). **B.** Illustration of *in vivo* engraftment of patient AMLs indicating bone marrow aspiration time points and treatment interval. Each arrow for CKS1i treatment refers to one day. **C.** Percentage of human CD45^+^ cells of total CD45^+^ cells in mouse bone marrow aspirations one week after chemotherapy (week 6). **D.** Kaplan-Meier survival curve and p value calculated for each individual PDX Control and CKS1i treated. Each data point represents one mouse. **E.***CKS1B* expression (relative to *GAPDH*) for patient AMLs tested under L-LTC-LIC conditions (N=7). **F.** Illustration of treatment time points, treatment and readout for *ex vivo* L-LTC-IC. **G.** Fold change L-LTC-IC frequency, CKS1i treatment versus control, after 7 weeks of co-culture. **H**. Individual 1/L-LTC-IC frequencies with upper and lower limits for each patient tested. A Student’s *t*-test was used to calculate significance of difference for all graphs unless otherwise stated. * p<0.05; **p<0.01; *** p< 0.001.

Whilst reducing leukemic blast count is the current backbone of clinical chemotherapeutic protocols, typically these approaches do not clear the most quiescent leukemic stem cells, the subset of cells at the origin of relapse *in vivo*(25,26). The observed effect on overall survival upon CKS1i treatment in *CKS1B*^low^ AMLs, without significant reduction of leukemic burden could indicate a direct effect of CKS1i on LSCs. We sought to assess this by using a leukemic-long-term culture initiating cell assay (L-LTC-IC)(27).

We selected seven patients with a range of *CKS1B* expression (Figure 2E) for *ex vivo* study (Figure 2F). After treatment, all surviving cells were re-plated at limiting dilution to assay the frequency of L-LTC-IC between control and CKS1i-treated cells. All patient samples showed significant reduction in L-LTC-IC frequency, regardless of bulk *CKS1B* expression at the start of the experiment (Figure 2G-H, Supp. Fig. 2B-C).

These data indicate that CKS1i can target *CKS1B*^high^ AML blasts, but more importantly is efficient at targeting the leukemic stem cell compartment. Considering these results, we hypothesized that the overall survival advantage *in vivo* of both *CKS1B*^*high*^ and *CKS1B*^*low*^ AMLs is likely due to depletion of LSCs, and therefore a smaller residual pool of these cells able to repopulate the AML overtime.

#### Normal and malignant hematopoietic cells have divergent responses to CKS1i

To understand the key mechanisms by which CKS1i kills AML, and what effects CKS1i has on normal hematopoietic cells, we performed mass spectrometry on *CKS1B*^high^ THP-1 AML cells and umbilical cord blood derived healthy CD34^+^ HSPCs (hereafter referred to as CD34^+^) pre- and post-treatment with CKS1i *in vitro* (1μM; Figure 3A).

**Figure 3.**
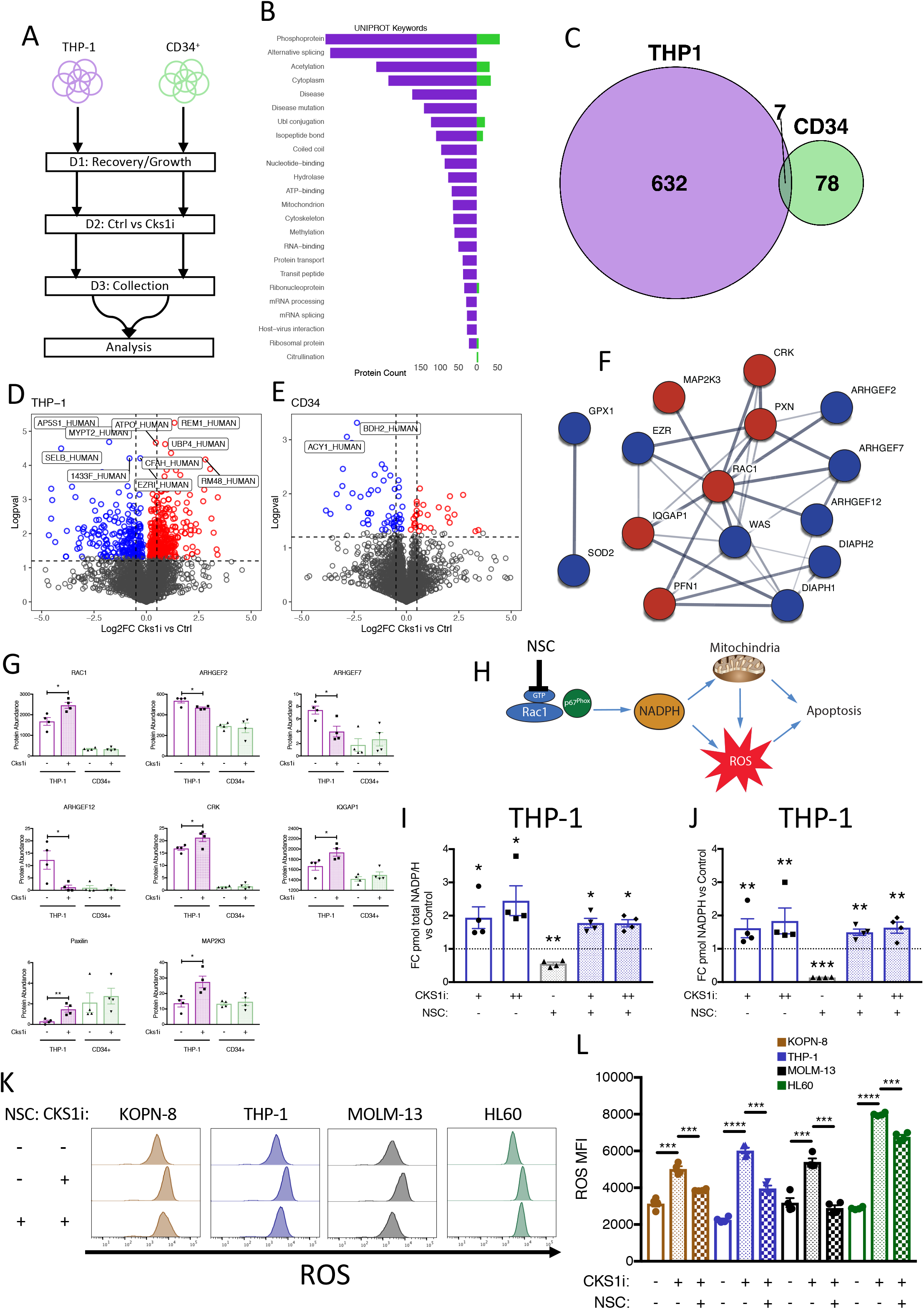
Divergent responses to CKS1i lead to RAC1-dependent induction of lethal ROS in AML. **A.** Workflow for timescale of cell preparation for mass spectrometry analysis. **B.** UNIPROT keywords for significantly enriched proteins in THP-1 (purple) and CD34^+^ (Green) cells in CKS1i treated conditions versus control. **C.** Venn diagram depicting overlap of differentially expressed proteins between THP-1 and CD34^+^ cells. Volcano plots for changes in protein abundance (CKS1i vs Control) in **D.** THP-1 and **E.** CD34^+^ cells. Red dots indicate significantly upregulated proteins, blue dots indicate significantly downregulated proteins (N=4 per cell line and treatment). **F.** String network analysis (Red = increased abundance, Blue = decreased abundance) and **G.** Protein abundance of key RAC1 pathway members differentially abundant in THP-1 but not CD34^+^ cells. **H.** Illustration of RAC1/NADPH/ROS pathway, indicating mode of action for RAC1 inhibitor NSC23766 (NSC). **I.** Fold change total NADP/NADPH and **J.** fold change NADPH in THP-1 cells treated with the indicated doses of CKS1i (+ = 1μM, ++ = 5μM) or NSC (+ = 0.1μM) versus control for 24 hours (N=4 per cell line and treatment). **K.** Representative flow plots and **L.** quantified mean fluorescence intensity of intracellular reactive oxygen species (ROS) in the indicated cell lines in response to CKS1i (+ = 1μM) and NSC (+ = 0.1μM) treatment (N=3 per cell line and treatment). A Student’s *t*-test was used to calculate significance of difference for all graphs. * p<0.05; **p<0.01; *** p< 0.001; **** p<0.0001.

Upon CKS1i treatment, the majority of differentially expressed proteins in THP-1 cells were phosphoproteins (387), which can be alternatively spliced at the RNA level (375) and also may be acetylated (257), demonstrating integration with post-translational modification (Figure 3B). CKS1i treatment induced divergent proteomic profiles in THP-1 and CD34^+^ cells, with differentially abundant proteins predominantly independent between healthy and malignant cells (Figure 3C-E).

Deeper pathway analysis of CKS1i-induced proteomic changes in THP-1 cells identified the upregulation of the small GTP-binding molecular switch protein RAC1, and its associated upstream and downstream regulators (Figure 3F-G), that are part of a pathway known to be involved in regulating cell growth and survival in response to a variety of external factors(28,29), and important for HSC homing and HSC/LSC niche interactions(30). Although multiple guanine nucleotide exchange factors (GEFs) are downregulated in CKS1i treated THP-1 cells, upregulation of the GTP loaders CRK and Paxilin indicate potential hyperactivity of RAC1, and in keeping with this, downstream MAP2K3 is also upregulated (Figure 3G).

The relative lack of RAC1 signalling pathway member changes in CD34^+^ proteomes are in keeping with RAC1-GTPase activity being upregulated in neoplastic transformation of hematopoiesis(30) and may identify a specific molecular switch in AML vulnerable to CKS1 inhibition.

#### CKS1i drives RAC1-dependent ROS accumulation in AML

The active GTP-bound form of RAC1 is known to bind to p67^Phox^ and catalyzes NADP/NADPH production. This in turn can lead to upregulation of intracellular reactive oxygen species (ROS), a process that has been shown to drive apoptosis and eliminate quiescent LSCs (Figure 3H)(31–33). In AML cell lines, CKS1i increased total NADP/NADPH levels in a dose dependent manner (Figure 3I, Supp. Fig. 3A), by increasing the total amount of NADPH compared to control (Figure 3J, Supp. Fig. 3B). The accumulation of NADPH in AML cell lines can partially be rescued by inhibiting RAC1 activity (Figure 3I-J, Supp. Fig. 3A-B), validating increased RAC1-GTP/p67^Phox^ activity upon CKS1 inhibition.

CKS1i-induced NADPH accumulation lead to significantly increased intracellular ROS in all AML cell lines at sublethal doses of CKS1i, regardless of *CKS1B* expression (Figure 3K-L). Reversal of this phenotype, by RAC1 inhibition, was most significant in *CKS1B*^high^ cell lines (Figure 3K-L). Finally, the reduction in AML cell line viability could be rescued by RAC1 inhibition, with *CKS1B*^high^ cell lines showing the strongest sensitivity (Supp. Fig. 3C-F). These data indicate that AML requires SCF^SKP2-CKS1^ functions to regulate RAC1 activity and maintain the fine balance of intracellular ROS, which are critical for LSC viability.

#### CKS1 inhibition protects normal hematopoiesis from stress

Since CKS1i is able to selectively kill *CKS1B*^*high*^ bulk AML, reduce the LSC compartment, and prolong AML xenograft survival regardless of leukemic reduction (Figure 2D), we hypothesized that normal hematopoiesis is spared by CKS1i treatment.

Proteomic analyses of CD34^+^ cells treated with CKS1i revealed a clear separation from THP-1 AML cells (Figure 3C-E). Key proteins differentially abundant in CD34^+^ cells and not THP-1 cells were integrated in three fundamental pathways in normal hematopoiesis: Wnt signalling, cell cycle and NFkB signalling (Supp. Fig. 4A-D). The changes in cell cycle are also consistent with previous reports of *Cks1*^*−/−*^ and CKS1i cellular phenotypes(10,34), and our previously reported suppression of Wnt signalling in *Cks1*^*−/−*^ mouse embryonic fibroblasts(9). The substantial change of proteins in these key pathways are hallmarks of growth suppression and differentiation, rather than an induction of cell death by CKS1i, and indeed, treatment of CD34^+^ cells *in vitro* with CKS1i resulted in reduced overall cell growth, with a blockage in cell cycle resulting in increased quiescence of CD34^+^ cells (Figure 4A-B).

**Figure 4.**
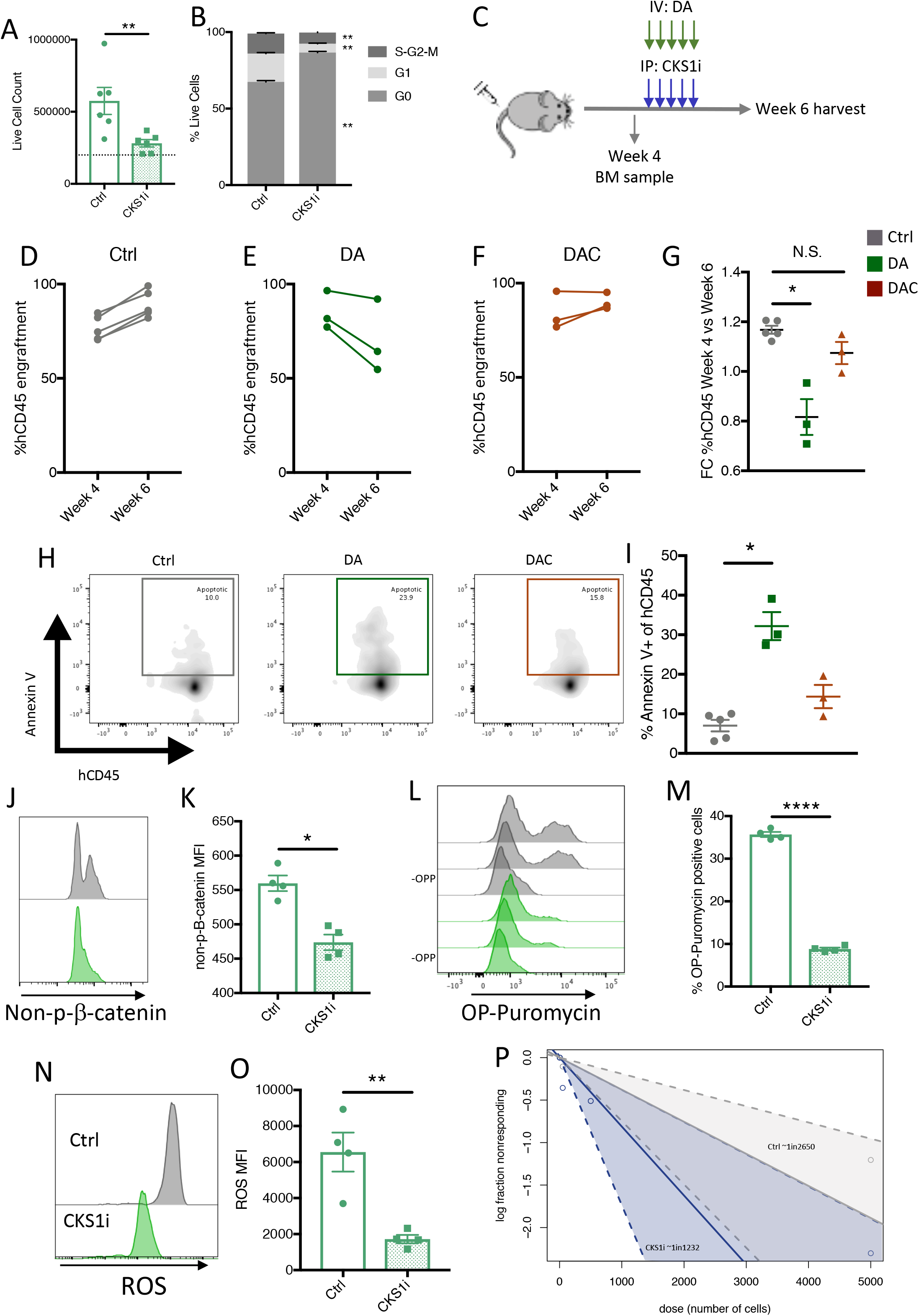
CKS1i protects normal hematopoietic cells from chemotherapeutic toxicity by suppressing key signalling pathways. **A.** Live cell count (N=6); **B.** Quantified cell cycle proportions of CD34^+^ cells grown for 48 hours in control conditions or treated with CKS1i. **C.** Illustration of CD34^+^ engraftment and chemotherapeutic treatment in NSG mice. Each dot corresponds to one mouse. Change in percentage human CD45^+^ of total CD45 at the indicated time points for **D.** Control, **E.** Doxorubicin/Cytarabine (DA) and **F.** Doxorubicin/Cytarabine plus CKS1i (DAC) treatments. **G.** Fold change % human CD45 cells week 4 to 6 for the indicated treatments. **H.** Representative flow plots and **I.**% total cells annexin V positive after 6 weeks *in vivo* for human CD45 cells with the indicated treatment conditions (Ctrl N=5, DA N=3, DAC N=3). **J.** Representative flow plots and **K.** quantified mean fluorescence intensity for non-phosphorylated β-catenin in CD34^+^ cells grown for 48 hours in control conditions or treated with CKS1i (N=4). **L.** Representative flow plots (including cells grown without OP-Puromycin; - OPP) and **M.**% total OP-Puromycin incorporation in CD34^+^ cells grown for 48 hours in control conditions or treated with CKS1i. OP-Puromycin was added 1hr prior to collection and fixation of cells (N=4). **N.** Representative flow plots and **O.** quantified mean fluorescence intensity of intracellular reactive oxygen species (ROS) in CD34^+^ cells grown for 48 hours in control conditions or treated with CKS1i (N=4). **P.** ELDA quantification plot for CD34^+^ cells grown in LTC-IC under control (grey) and CKS1i (blue) treated conditions. Linear model and confidence interval are shown with estimated stem cell frequency indicated. A Student’s *t*-test was used to calculate significance of difference unless otherwise stated. * p<0.05; **p<0.01; *** p< 0.001; **** p<0.0001.

Considering that suppression of the cell cycle in CD34^+^ by CKS1i would reduce the potential for integration of nucleotide analogues and the requirement for topoisomerase activity, we hypothesized that CKS1i treatment may be “chemo-protective” for normal hematopoietic cells. To test this, we engrafted normal CD34^+^ cells in NSG mice and treated mice with the clinical chemotherapy protocol of cytarabine plus doxorubicin (as has been previously published; 5+3(35)), with or without CKS1i (Figure 4C). Human CD45^+^ bone marrow engraftment increased in untreated control mice between weeks 4 and 6 (Figure 4D). Treatment at week 4, with doxorubicin/cytarabine (DA) significantly reduced bone marrow engraftment by week 6, but addition of CKS1i (DAC) was able to rescue this effect, returning engraftment to comparable levels to control (Figure 4E-G). Better engraftment at week 6 was complemented by a reduction in apoptotic human CD45 cells in the bone marrow of recipient mice (Figure 4H-I), indicating that CKS1i treatment prevents DA-induced cell death in normal hematopoietic cells.

Mechanistically, mass cytometric analysis of CKS1i-treated CD34^+^ cells revealed a conserved reduction in key active signalling and transcriptional components that we recently reported to be important for proliferation and differentiation of HSPCs during *in vitro* amplification, including NFkB, PU.1, CREB and mTOR (Supp. Fig. 5)(36), further indicating an overall block in normal hematopoietic stem cell function. Indeed, it is important to note that whilst classical cell cycle phosphorylation marks (e.g. CDK1^pT14/Y15^) are reduced, the protein levels of differentiation regulators are also reduced (e.g. CREB, PU.1), indicating a potential block in differentiation as well as growth (Supp. Fig. 5). Additionally, fewer cells have active β-catenin, indicating that the Wnt pathway – a fundamental pathway that requires a tight balance for normal hematopoiesis to proceed(37) – is also suppressed (Figure 4J-K). Further suppression of markers of metabolically active cells (e.g. mTOR^pS2448^), inflammatory responses (e.g. NFkB^pS529^), and suppression of translation machinery in our mass spectrometry analyses, lead to a reduction in protein production in CKS1i treated CD34^+^ cells (Figure 4L-M). Together, these signalling pathways are fundamentally linked to the control of ROS accumulation in HSCs, with high activity in these pathways leading to high intracellular ROS, reduced cell viability and reduced stem cell potential(38). In agreement, CKS1i treatment reduced intracellular ROS in cultured CD34^+^ cells (Figure 4N-O) and *ex vivo* CKS1i treatment increased LTC-IC frequency by more than two-fold (1 in 1232 vs 1 in 2650, Figure 4P).

Together, these data indicate that the temporary suppression of HSPC activity conferred by CKS1i leads to suppression of cell growth, protection from general metabolic stress, improved stem cell functionality and overall increased healthy hematopoietic capacity during induction chemotherapy.

Combining CKS1 inhibition with induction chemotherapy protects normal hematopoiesis, reduces leukemic stem cells and improves overall survival

The current frontline chemotherapeutic protocols for AML include the use of DA as induction therapy agents. To test the potential for combining classical DA chemotherapy with CKS1i *in vivo*, we transplanted NSG mice with primary AML samples with varying *CKS1B* expression (Figure 5A-B), and after stratifying for engraftment at week 4, we treated the mice with either DA or DAC. One-week post chemotherapy, xenografts showed strong reduction in leukemic burden in both DA and DAC treatment for all AMLs, regardless of *CKS1B* expression, indicating that CKS1i does not interfere with normal chemotherapeutic killing (Figure 5C). At the same time point, resident murine CD45 cells co-extracted from aspirated tibias had significantly higher colony forming potential upon the addition of CKS1i compared to untreated mice and DA treated mice (Figure 5D), indicating that CKS1i treatment can both selectively reduce AML, whilst protecting normal hematopoiesis from chemotherapeutic toxicity. Overall, all xenografts showed a trend towards improved overall survival with DA treatment, and in all patients this was significantly improved by the addition of CKS1i, in line with *CKS1B* expression of these patients (Figure 5E-H).

**Figure 5.**
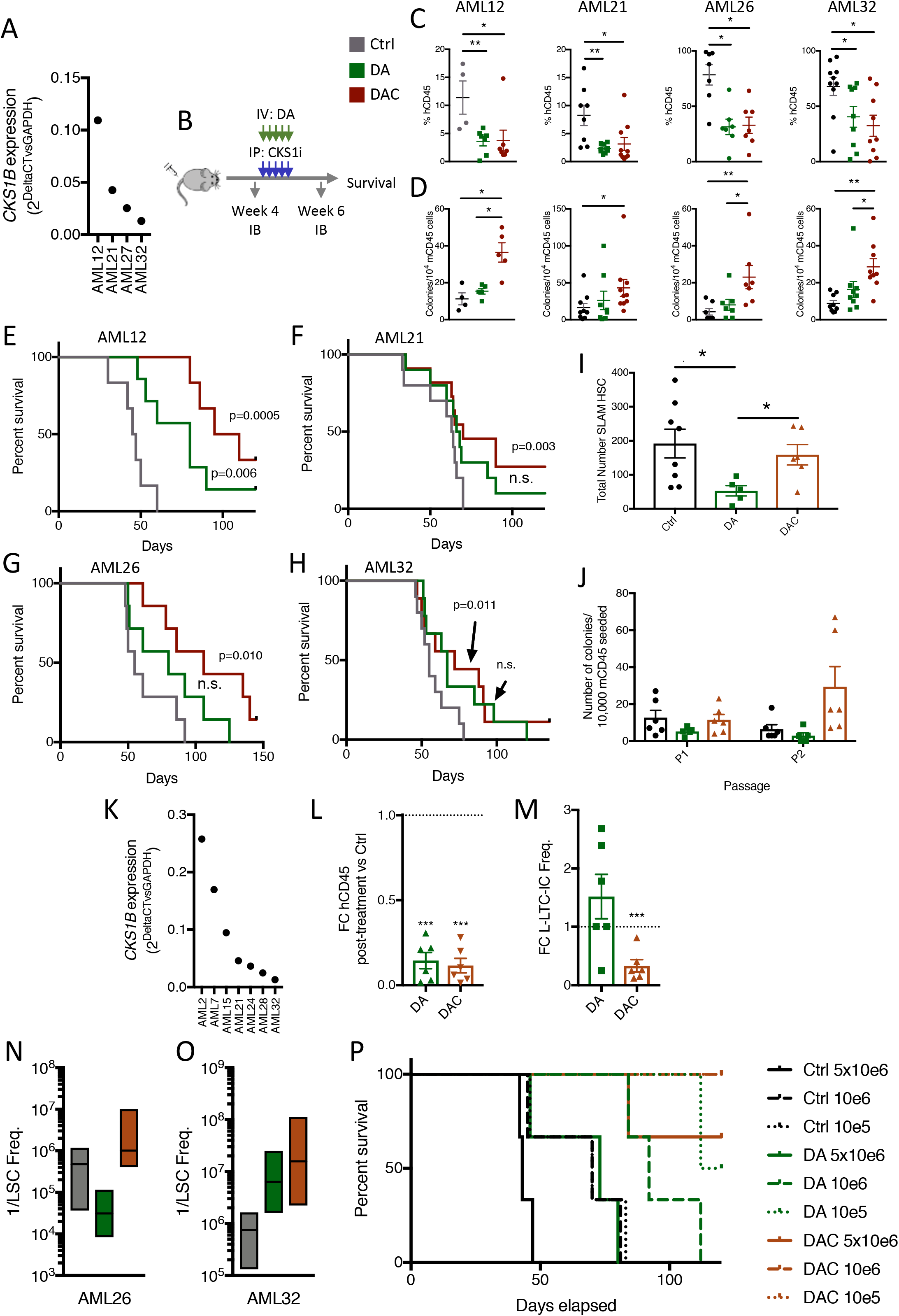
Combination induction chemotherapy and CKS1i reduces AML burden and leukemic stem cell potential whilst protecting resident hematopoietic cells. **A.** *CKS1B* expression (relative to *GAPDH*) for patient AMLs tested *in vivo* (N=4). **B.** Illustration of *in vivo* engraftment of patient AMLs indicating bone marrow aspiration time points and treatment interval. Each arrow for DA and CKS1i treatment refers to one day. **C.** Percentage of human CD45^+^ cells of total CD45^+^ cells in mouse bone marrow aspirations one week after chemotherapy (week 6). **D.** Colony forming units per 10,000 mouse CD45^+^ cells seeded from week 6 bone marrow aspirations. **E-H.** Kaplan-Meier survival curve and p value calculated for each individual PDX Control and treated mice. Each data point represents one mouse. **I.** Total number of murine Long-term HSCs obtained from bone marrow of mice at the final survival time point (Ctrl N=8, DA N=5, DAC N=5). **J.** Serial colony forming units per 10,000 mouse CD45^+^ cells obtained from BM of mice at the final survival time point (Ctrl N=6, DA N=5, DAC N=6). **K.***CKS1B* expression (relative to *GAPDH*) for patient AMLs tested in L-LTC-IC (N=7). **L.** Fold change live human CD45^+^ cells, indicated treatments versus control, after two weeks of co-culture. **M.** Fold change of L-LTC-IC frequency, indicated treatment versus control, after 7 weeks of co-culture. **N-O.** LSC frequency in secondary transplanted mice injected with AML26 **N.** and AML32 **O.** at limiting dilution 6 weeks post-transplantation. **P.** Kaplan-Meier survival curve for all AML32 secondary mice up to 120 days. A Student’s *t*-test was used to calculate significance of difference for all graphs unless otherwise stated. * p<0.05; **p<0.01; *** p< 0.001; **** p<0.0001.

Examination of the normal hematopoietic compartment of xenografted mice at the end point of survival, revealed a severe reduction in total number of long-term HSCs (SLAM/LT-HSC) in the DA treated group, whereas addition of CKS1i to DA abolished this effect (Figure 5I). In addition, serial colony forming ability of normal murine hematopoietic cells was improved in DAC conditions, indicating that rescued LT-HSCs are still functional (Figure 5J).

We and others have well documented the refractory nature of LSCs to induction chemotherapy(24,39), and here we set out to investigate the potential conflict or beneficial contribution between DA enrichment of LSCs and CKS1i depletion of LSCs. In *ex vivo* conditions, *CKS1B* high and low AMLs (Figure 5K) showed significant reduction in total cell number one week after DA or DAC treatment (Figure 5L), yet whilst DA treatment enriched for L-LTC-IC frequency in three of the six patients, CKS1i markedly reduced L-LTC-IC frequency in all patients (DAC; Figure 5M & Supp. Fig 6A-B).

Finally, to test the reduction in LSC frequency conferred by CKS1i *in vivo*, we engrafted AML cells obtained from AML26 and AML32 (which had the smallest improvement in overall survival after chemotherapy) in secondary recipients in a limiting dilution manner. Whilst control AMLs retained strong LSC frequency and show robust engraftment after 6 weeks, this was increased by DA treatment in AML26 and was notably reduced in AML32 (Figure 5N-O, Supp. Fig. 7A-B). The addition of CKS1i counteracted the effect of DA by decreasing the LSC frequency in AML26 and further reducing LSC frequency in AML32 compared to control mice, demonstrating strong reduction in LSCs after CKS1i treatment (Figure 5N-O, Supp. Fig. 7A-B). Overall survival was tested for AML32, with DA-AML mice surviving significantly longer than controls, and DAC-AML mice demonstrating a further improvement, with no overt signs of sickness at 120 days in all but one case at the highest cell dose (Figure 5P).

Together, these data indicate that inhibition of the SCF^SKP2-CKS1^ complex has dual roles, reducing the viability of AML, and importantly of the LSC fraction, whilst protecting normal HSPCs from chemotherapeutic stress.

## Discussion

In this study, we demonstrate that proteostatic targeting of the SCF^SKP2-CKS1^ E3 ubiquitin ligase complex selectively eliminates *CKS1B*^high^ AML blasts and reduced the LSCs compartment of *CKS1B*^high & low^ AMLs, while sparing normal hematopoietic cells from chemotherapeutic toxicity.

Poor risk AML is a heterogeneous group that includes patients with different cytogenetic abnormalities, very limited treatment options and extremely low overall survival rates(1,40), even accounting for newer therapies (Venetoclax plus Azacitidine)(2,3). The selective reduction in viability of *CKS1B*^high^ AML blasts by CKS1 inhibition (Figure 1D) indicate that, whilst *CKS1B* is not predictive of overall survival at the RNA level, proteostatic vulnerabilities exist in AML and can be identified through better understanding of leukemic proteomes. While gene expression profiles, particularly those with single cell resolution, are improving our understanding of AML heterogeneity, the origins of leukemic relapse and revealing new clinical targets(25,26), the role of proteostasis has been comparatively understudied(41,42).

*In vivo*, AML blasts and LSCs are relatively quiescent compared to AML cell lines and patient samples screened *in vitro*, so the consistently high sensitivity of *CKS1B*^*high*^ AMLs observed *in vivo* (Figure 2C) and the reduction of LSC frequency regardless of the bulk *CKS1B* expression (Figure 2E-H), demonstrates a role for the SCF^SKP2-CKS1^ complex beyond targeting highly proliferative cells. Therefore, the role of CKS1 likely reaches further than cell cycle regulation, as has been previously reported(10,43). Here we provide evidence for CKS1 regulating RAC1/NADPH/ROS signalling (Figure 3H), a fundamental pathway involved in amplifying extrinsic and intrinsic signals in normal hematopoiesis and AML(4,44). The balance of intracellular ROS in normal and malignant hematopoietic cells has been of great interest in recent years(33,38), and changes in mitochondrial functions due to *RAS* mutations and nicotinamide-NAD metabolism underline the critical role for this pathway in primary patient resistance to Venetoclax(4,5). The induction of ROS in AML cell lines upon CKS1 inhibition, regardless of *CKS1B* expression, demonstrates that the balance of CKS1-dependent protein degradation is key to maintaining stress responses in AML. This, together with LSCs requiring low ROS to maintain their stem cell potential, would explain the strong reduction in LSC frequency conferred by CKS1i in primary patient AML (Figure 2G-H & 5M-O).

The effect of CKS1i on normal hematopoiesis is clearly separate to AML (Figure 3C). Indeed, cell cycle blockage was suggested by the relatively few protein changes induced by CKS1i in CD34^+^ but not AML cells. This is highly beneficial, as patients treated with induction chemotherapy, which targets cycling cells, suffer from severe toxicity and cytopenia upon treatment. Classical induction chemotherapy is known to reduce the pool of hematopoietic progenitors, whilst quiescent HSCs are refractory to treatment, but ultimately undergo senescence(7,45). We found that cell cycle arrest of HSPCs by CKS1i could prevent DA reduction of normal cells *in vivo* (Figure 4D-G), and in the context of AML could rescue the reduction in HSCs induced by chemotherapy (Figure 5I). Importantly, CKS1i treatment also induced changes in fundamental HSPC signalling pathways known to be involved in stem cell potency and response to stress. Indeed, we have previously shown that activating NFκB signalling can reduce intracellular ROS and improve HSC outgrowth *in vitro*(36). The overall suppression of key growth and activation cellular markers lead to an opposite phenotype to that seen in AML cells, with a reduction in intracellular ROS and an increase in normal HSC frequency (Figure 4N-P). These divergent phenotypes between normal and malignant hematopoietic cells indicate that CKS1-dependent protein degradation is required for the growth of normal cells and that CKS1 suppression pushes cells towards quiescence while retaining stem cell functions. On the other hand, inhibiting CKS1-dependent protein degradation in AML can lead to incorrect regulation of signalling, ultimately causing cellular toxicity, as shown before for more broader regulators of the SCF complex(46).

The backbone of clinical chemotherapeutic protocols has largely remained unchanged over the last 20 years, with induction chemotherapy reducing AML blasts to prolong survival(47). Whilst reduction of AML burden is beneficial, classical chemotherapy actually maintains or even enriches for LSCs and results in relapsed AML, often with increased mutational burden(48). The addition of CKS1i to DA results in a significant reduction in LSC frequency. Considering that resistant LSCs can be traced as the origin of relapse(25,26), and constitute a key route to chemoresistance in AML, CKS1 inhibition has excellent promise for eradicating these cells.

The inhibition of CKS1-dependent protein degradation holds excellent promise for AML therapy, both as a targeted agent towards *CKS1B*^*high*^ AML, and in combination with induction chemotherapy where protection of healthy cells is key. Reports of *CKS1B* overexpression correlating with outcome in other solid cancer types(20,22,49) and novel ways to modulate CKS1 function(50), together with our findings of a dual role for CKS1 inhibition in AML and the development of more clinically ready molecules to target CKS1(51), indicate that proteostatic targeting, through the CKS1/CKS2 axis, holds much hope for future cancer therapy.

## Methods (online unlimited)

### Primary AML and UCB samples

AML samples were obtained after informed consent at St Bartholomew’s Hospital (London, U.K.) at the time of diagnosis as part of the Bart’s Cancer Institute Poor-Risk AML consortium. Full details of patient information are provided in Supplementary Table 1. Live mononuclear cells (MNCs) were isolated by density centrifugation using Ficoll-Paque (GE healthcare). Prior to culture or xenotransplantation AML cells were depleted for T-cells using the Easysep T-cell depletion kit (StemCell Technologies). Umbilical Cord Blood (UCB) was obtained from full term donors after informed consent at the Royal London Hospital (London, U.K.). MNCs were isolated by density centrifugation using Ficoll-Paque (GE healthcare). Cells were selected for CD34^+^ using the Easysep CD34^+^ enrichment kit (StemCell Technologies). Purity was confirmed by flow cytometry.

### Drug sensitivity and resistance testing (DSRT)

Single drug DSRT was performed as described previously(52). In brief, 35 different compounds, each with 7 different concentrations (Supp. Table. 2), were pre-plated using an acoustic liquid handling Echo 550 (Labcyte) to 384-well plates. Drug plate well annotations and drug concentrations are presented in Supp. Table. 3. Primary AML cells were suspended in conditioned medium (RPMI 1640 supplemented with 10% foetal bovine serum, 2mM L-glutamine, penicillin-100U/ml, streptomucing-100ug/ml and 12.5% conditioned medium from HS-5 human bone marrow stromal cells), DNase I treated for 4h (Promega), filtered through a 70μm cell strainer (Fisher Scientific) to remove possible cell clumps, and viable cells were counted. Pre-plated compounds in each 384-well plate were dissolved in 5ul of conditioned medium using a MultiDrop Combi peristaltic dispenser (Thermo Scientific) and shaken for 5 minutes to dissolve the compounds. AML cells were plated at 5,000 cells/well in 20ul, leading to a final volume of 25ul/well. Plates were gently shaken for 5 minutes to mix the cells with the compounds and incubated for 72 hours at 37C, 5% CO_2_.

Cell viability was measured using the CellTiter-Glo assay (Promega) with a PHERAstar microplate reader (BMG-labtech). Data was normalised to negative (DMSO only) and positive control wells (100uM benzethonium chloride) and dose response curves calculated.

*Ex vivo* drug sensitivity of AML cells to the tested drugs was calculated using a drug sensitivity score (DSS), a modified form of the area under the inhibition curve calculation that integrates multiple dose response parameters for each of the tested drugs, as previously described(53).

#### AML cell line, UCB CD34^+^ and MS-5 culture

All AML cell lines and MS-5 stromal cells were originally obtained from the ATCC and maintained by the Francis Crick Cell Services. All AML cell lines were cultured in RMPI 1640, 10% heat-inactivated FBS and 1% penicillin/streptomycin (Life Technologies) at 37C, 5% CO_2_. Umbilical cord blood CD34^+^ cells were cultured in StemSpan SFEMMII (StemCell Technologies) supplemented with Human SCF (150ng/ml), Human FLT3 ligand (150ng/ml) and Human TPO (20ng/ml; all Peprotech) at 2×10^5^ cells/ml at 37C, 5% CO_2_. For relative viability, apoptosis and IC_50_ calculations cell lines were seeded in 96 well plates at a concentration of 2×10^5^ cells/ml with the indicated dose of drug. Measurements of viability (% reduction O_2_) or apoptosis (Annexin V positivity) were taken at 48 hours post treatment. MS-5 stromal cells were cultured in IMDM, 10% heat-inactivated FBS and 2% penicillin/streptomycin (Life Technologies) at 37C, 5% CO_2_.

#### Publicly available datasets

*CKS1B* expression in normal and malignant hematopoiesis was obtained through Bloodspot.eu. Overall survival and stratification for *CKS1B* expression was calculated from data obtained from The Cancer Genome Atlas (TCGA). AML cell line RNA sequencing data was obtained from the EBI Expression Atlas (RNA-seq of 934 Human cancer cell lines from the Cancer Cell Line Encyclopedia).

#### Leukemic/Normal Long-term culture initiating cell (L-LTC-IC) assay

These experiments were performed as originally published by our group(27). For all co-culture experiments, MS-5 stromal cells were seeded two days prior to AML/UCB cell addition at 4×10^5^ cells/ml to reach confluence at the time of irradiation. One day prior to AML/UCB addition, MS-5 stromal cells were irradiated with 7Gy and culture media was exchanged. On the day of starting co-culture, AML cells were plated at 2×10^5^ cells/ml in meylocult H5100 (StemCell Technologies) supplemented with IL-3, G-CSF and TPO (all 20ng/ml; Peprotech). UCB cells were plated at 2×10^5^ cells/ml in myelocult H5100 (StemCell Technologies). Half media changes were performed once per week without disrupting the feeder layer. At the start of week two, indicated drug treatments were added at 2x concentration in the half media change.

For LTC-CAFC assays, all cells were harvested at day 14 and sorted for live hCD45^+^mSca-1^−^ cells. Resulting cells were seeded in co-culture with fresh MS-5 stromal cells in a 96 well plate in a limiting dilution range (200,000 to 1,000) in 10 replicates and cultured for a further 5 weeks. At the end of the co-culture period cobblestone area forming cells were scored and L-LTC-IC frequency was calculated using the ELDA (Extreme Limiting Dilution Analysis) function in the Statmod R package.

For LTC-IC assays, media was continuous changed each week until week five Cultures were harvested and sorted for live hCD45^+^mSca-1^−^ cells. Resulting cells were seeded in co-culture with fresh MS-5 stromal cells in a 96 well plate in a limiting dilution range (10,000 to 100) in 10 replicates and cultured for a further three weeks. At week eight, myelocult H5100 was replaced with Methocult methycellulose (StemCell Technologies H4434) for a further two weeks, after which wells were scored for colony-forming units and LTC-IC frequency was calculated using the ELDA (Extreme Limiting Dilution Analysis) function in the Statmod R package.

#### Patient derived xenografts (PDX) and *in vivo* drug treatment

Primary AML samples (1×10^6^ – 5×10^6^ cells total) or UCB-CD34^+^ (5×10^4^ cells total) were injected intravenously (I.V.) into unconditioned 10-12 week old female or male NOD-SCID IL2Rynull (NSG) mice (The Jackson laboratory). After 4 weeks engraftment was assessed by bone marrow aspiration from long bones whilst mice were under isoflurane anaesthesia. Mice were stratified according to engraftment and sex and assigned to treatment and control groups accordingly. Mice were treated as indicated with 10mg/kg CKS1i (Skp2-Cks1 E3 ligase inhibitor, Merck Millipore) intraperitoneal injection (I.P.) for 5 days, DA (doxorubicin/cytarabine, 1.5mg/kg/10mg/kg respectively, Sigma Aldrich), doxorubicin on days 1-3, cytarabine on days 1-5 co-injected I.V.(35). Mice were scored for engraftment over the experimental course by bone marrow aspiration and for overall survival according to U.K. home office license protocols and following CRUK guidance (>20% peak body weight loss, overt signs of sickness/mortality).

#### AML cell line *in vivo* experimentation

AML cell lines were transduced with GFP-Luciferase containing vectors as per our previous reports(24). For both cell lines (THP-1 and HL60) 2×10^6^ cells were injected I.V. into unconditioned 10-12 weeks old female or male NOD-SCID IL2Rynull (NSG) mice (The Jackson laboratory). After 7 days engraftment was assessed by bioluminescene imaging. Isofluorane anaesthetised mice were imaged 5-10 minutes post D-luciferin injection I.P. (15mg.kg; Caliper life sciences) using the Xenogen IVIS imaging system. Photons emitted were expressed as Flux (photons/s/cm^2^), and quantified and analysed using “living image” software (Caliper life sciences).

#### RNA extraction, reverse transcription and real time quantitative PCR (RT-qPCR)

Total RNA was isolated from patient samples after thawing, density centrifugation and T-cell depletion, using a RNeasy mini kit (Qiagen). Resulting RNA was reverse transcribed to produce cDNA using the Superscript III reverse transcriptase kit (Thermo Scientific) with oligoDT_20_ primers (Sigma Aldrich). RT-qPCR experiments were performed with an ABI-7500 FAST Thermal Cycler (Applied Biosystems) using SYBR Green (Thermo Scientific). RNA abundance was quantified by the Comparative CT method with two independent control genes (*GAPDH* and *B-ACTIN*, *GAPDH* presented). The CT values used for each patient sample were the result of three technical triplicates. Primers are described in the resources table.

#### Mass Spectrometry

THP-1 AML cell lines and UCB CD34^+^ cells were cultured as per culture and drug treatment above. Cells were recovered for 24 hours in their respective media followed by sub-lethal AML doses of CKS1i (1μM) for 12 hours. All cells were retrieved from wells, washed three times in ice-cold PBS and snap frozen in liquid nitrogen as dry pellets.

Cell pellets were lysed in 100 μL of urea buffer (8 M urea in 20 mM HEPES, pH: 8.0), lysates were further homogenized by sonication (30 cycles of 30s on 30s off; Diagenode Bioruptor® Plus) and insoluble material was removed by centrifugation. Protein amount was quantified using BCA (Thermo Fisher Scientific). Then, 100 and 20 μg of protein for THP-1 and CD34^+^ samples, respectively, were diluted in urea buffer to a final volume of 300 μL and subjected to cysteine alkylation using sequential incubation with 10 mM dithiothreitol (DDT) and 16.6 mM iodoacetamide (IAM) for 1 h and 30 min, respectively, at 25 °C with agitation. Trypsin beads (50% slurry of TLCK-trypsin; Thermo-Fisher Scientific; Cat. #20230) were equilibrated with 3 washes with 20 mM HEPES (pH 8.0), the urea concentration in the protein suspensions was reduced to 2 M by the addition of 900 μL of 20 mM HEPES (pH 8.0), 100 μL of equilibrated trypsin beads were added and samples were incubated overnight at 37°C. Trypsin beads were removed by centrifugation (2000 xg at 5°C for 5 min) and the resulting peptide solutions were desalted using carbon C18 spin tips (Glygen; Cat. # TT2MC18). Briefly, spin tips were activated twice with 200 μL of Elution Solution (70% ACN, 0.1% TFA) and equilibrated twice with 200 μL of Wash Solution (1% ACN, 0.1% TFA). Samples were loaded and spin tips were washed twice with 200 μL of Wash Solution. Peptides were eluted into fresh tubes from the spin tips with 4 times with 50 μl of Elution Solution. In each of the desalting steps, spin tips were centrifuged at 1,500xg at 5C for 3 min. Finally, samples were dried in a SpeedVac and peptide pellets were stored at −80°C.

For mass spectrometry identification and quantification of proteins, samples were run twice in a LC-MS/MS platform. Briefly, peptide pellets were resuspended in 100 μL and 20 μL of reconstitution buffer (20 fmol/μL enolase in 3% ACN, 0.1% TFA) for THP-1 and CD34^+^ samples, respectively. Then, 2 μL were loaded onto an LC-MS/MS system consisting of a Dionex UltiMate 3000 RSLC coupled to a Q Exactive™ Plus Orbitrap Mass Spectrometer (Thermo Fisher Scientific) through an EASY-Spray source (Cat. # ES081, Thermo Fisher Scientific). Mobile phases for the chromatographic separation of the peptides consisted in Solvent A (3% ACN: 0.1% FA) and Solvent B (99.9% ACN; 0.1% FA). Peptides were loaded in a micro-pre-column (Acclaim™ PepMap™ 100 C18 LC; Cat. # 160454, Thermo Fisher Scientific) and separated in an analytical column (Acclaim™ PepMap™ 100 C18 LC; Cat. # 164569, Thermo Fisher Scientific) using a gradient running from 3% to 23% over 120 min. The UPLC system delivered a flow of 2 μL/min (loading) and 300 nL/min (gradient elution). The Q-Exactive Plus operated a duty cycle of 2.1s. Thus, it acquired full scan survey spectra (m/z 375–1500) with a 70,000 FWHM resolution followed by data-dependent acquisition in which the 15 most intense ions were selected for HCD (higher energy collisional dissociation) and MS/MS scanning (200– 2000 m/z) with a resolution of 17,500 FWHM. A dynamic exclusion period of 30s was enabled with a m/z window of ±10 ppms.

Peptide identification from MS data was automated using a Mascot Daemon 2.5.0 workflow in which Mascot Distiller v2.5.1.0 generated peak list files (MGFs) from RAW data and the Mascot search engine (v2.5) matched the MS/MS data stored in the MGF files to peptides using the SwissProt Database (SwissProt_2016Oct.fasta). Searches had a FDR of ~1% and allowed 2 trypsin missed cleavages, mass tolerance of ±10 ppm for the MS scans and ±25 mmu for the MS/MS scans, carbamidomethyl Cys as a fixed modification and PyroGlu on N-terminal Gln and oxidation of Met as variable modifications. Identified peptides were quantified using Pescal software in a label free procedure based on extracted ion chromatograms (XICs). Thus, the software constructed XICs for all the peptides identified across all samples with mass and retention time windows of ±7 ppm and ±2 min, respectively and calculated the area under the peak. Individual peptide intensity values in each sample were normalized to the sum of the intensity values of all the peptides quantified in that sample. Data points not quantified were given a peptide intensity value equal to the minimum intensity value quantified in the sample divided by 10. Protein intensity values were calculated by adding the individual normalized intensities of all the peptides comprised in a protein and values of 2 technical replicates per sample were averaged. Protein score values were expressed as the maximum Mascot protein score value obtained across samples. The mass spectrometry proteomics data have been deposited to the ProteomeXchange Consortium via the PRIDE partner repository (PXD022754 and 10.6019/PXD022754).

#### Flow Cytometry, apoptosis and cell cycle assays

Flow cytometry analysis was performed using a BD Fortessa flow cytometer (BD biosciences). Cells were prepared by washing in PBS + 1% FBS three times before staining in the same media with the indicated cell surface antibodies (resources table) for 1 hour at 4C. For apoptosis assays, cells were incubated with annexin V binding buffer in addition to the washing media (BD biosciences), washed three times in PBS + 1% FBS + 1x annexin V binding buffer and incubated with 0.1μg/ml DAPI prior to flow cytometry analysis. For cell cycle analysis, cells were washed three times in PBS + 1% FBS and fixed in BD fix/perm buffer (BD biosciences) for 20 minutes at room temperature. Cells were washed three times in BD perm/wash buffer (BD biosciences) and incubated with anti-Ki67 antibody for 4 hours at 4C. Cells were washed three times in BD perm/wash buffer and 0.5 μg/ml DAPI was added for 15 minutes prior to analysis. For all flow cytometry, cells were initially identified based on forward and side scatter.

#### Viability assays

Relative cell viability was assessed by % reduction O_2_ in culture wells using the Alamar blue cell viability reagent (Life Technologies). Cells were seeded in 96 well plates at 2×10^5^ cells/ml and the indicated dose of drugs were added on top and incubated for 48 hours. Alamar blue reagent was added on top of cells and they were continued to be incubated for 4 hours under the same conditions (37C, 5% CO_2_). Plates were read on a spectramax plate reader (Biostars) at 570nm and 600nm and % reduction O_2_ was calculated as per the manufacturer’s instructions.

#### NADP/NADPH assays

Total NADP^+^ and NADP/H were measured using the NADP/NADPH colorimetric assay kit (Abcam). AML cell lines were seeded at 2×10^5^ cells/ml one day prior to treatment with the indicated drugs (day 0). The following day (day 1), the indicated concentration of drugs were added to culture wells. The next day (day 2) all cells were collected from the wells and washed three times in ice-cold PBS. Cells were lysed in NADP/NADPH extraction buffer by performing two freeze/thaw cycles (20 mins on dry ice followed by 10 mins at room temperature). Lysates were centrifuged at 13,000g for 10minutes and the supernatant was retained. Lysate supernatant was split in half, with one half remaining on ice and the other half incubated at 60C for 30mins to remove NADP^+^. Total NADP/H (NADPt) and NADPH only lysates were run in 96 well plates with freshly made standards as per the manufacturers’ instructions. NADP/NADPH ratio was calculated as (NADPt-NADPH)/NADPH.

#### Intracellular ROS staining

Intracellular reactive oxygen species were assayed using the CellRox deep red reagent (Life Technologies). AML cell lines were seeded at 2×10^5^ cells/ml one day prior to treatment with the indicated drugs (day 0). The following day (day 1), the indicated concentration of drugs were added to culture wells. The next day (day 2), CellRox deep red was added to each well at a final concentration of 5uM and verapamil was added at a final concentration of 50 μM. Cells were continued to be incubated in the same conditions (37C, 5% CO_2_) for 1hr. After incubation, cells were collected from wells and washed three times in PBS + 1%FBS + 50 μM verapamil and finally resuspended in PBS + 1% FBS + 50 μM verapamil + DAPI (0.1μg/ml) before analysis on a BD Fortessa FACS analyser.

#### Mass Cytometry

CyTOF preparation and analysis was carried out as per our previous publication(36). Cultured cells were washed in ice-cold PBS three times and incubated with 5uM Cisplatin (Fluidigm) to mark dead cells. Cells were washed three times in ice-cold PBS and fixed in 1.6% formaldehyde (Sigma Aldrich). Fixed cells were surface stained with the relevant antibodies (resources table) for two hours at room temperature followed by three washes with PBS. Cells were permeabilised in 1ml Perm buffer III (BD biosciences) on ice for 30mins, washed three times in ice-cold PBS and incubated with the relevant intracellular antibodies (resources table) overnight at 4C with gentle rotation. Resulting cells were wash three times in ice-cold PBS and stained with 100nM Iridium in PBS + 0.1% Saponin (Riedel-de Haen) overnight before analysis on a Helios Mass Cytometer (Fluidigm).

#### Protein translation assays

Protein translation was measured using the OP-Puromycin protein translation kit (Life Technologies). AML cell lines were seeded at 2×10^5^ cells/ml one day prior to treatment with the indicated drugs (day 0). The following day (day 1), the indicated concentration of drugs were added to culture wells. The next day (day 2), 10 μM OP-Puromycin was added to culture wells for one hour under culture conditions (37C, 5% CO_2_). Cells were washed three times in ice-cold PBS and fixed in 4% paraformaldehyde (Sigma Aldrich) at room temperature for 15 mins in the dark. Cells were washed three times in PBS and permeabilised in PBS + 0.5% Triton X-100 (Sigma Aldrich) for 15 mins. Cells were washed twice in Click-IT reaction buffer wash solution and stained as per the manufacturer’s instructions (Life Technologies). Abundance of OP-Puromycin was assessed using flow cytometry.

#### Colony forming units

For resident mouse hematopoietic cell response to 5-FU’, CKS1i, DA and DAC, colony forming ability was assessed in methylcellulose (StemCell Technologies M3434-GF). 10^4^ mCD45^+^ cells were sorted from PDX mice at the indicated points and seeded in methylcellulose and scored to colony forming units after 7 days. Cultures were dissolved in PBS, counted and 10^4^ cells were re-seeded for passage 2 and passage 3.

#### Statistics and data interpretation

Results shown are +/-SEM unless otherwise indicated. To compare treatment versus control in all *in vitro* and *in vivo* experiments, a Student’s *t-*test was used as indicated in the figure legend with N number indicated. For all comparisons, unpaired *t*-tests were undertaken unless otherwise indicated. All repeat samples presented are from biological replicates of distinct samples/xenotransplantations.

Survival analyses were carried out using the “survminer” package on R to calculate significance between Kaplan-Meier curves and Hazard ratios. Kaplan Meier graphs were plotted using Graphpad Prism.

Correlation analyses were carried out using the “performance analytics” and “corrplot” packages in R. Multiple DSS comparisons with *CKS1B* expression were carried out with pairwise complete observations using Spearman, Pearson and Kendall correlation coefficients. Individual correlations for *CKS1B* vs DSS or IC_50_ were plotted using Graphpad Prism.

Stem cell frequency was calculated using the extreme limiting dilution analysis (ELDA) function in the “statmod” R package(54).

Pathway analysis and enrichment was run through MetaCore (genego.com) and network interactions produced on String (string-db.org)

## Acknowledgements

We would like to acknowledge the Francis Crick core flow cytometry and biological research facility STPs. The Francis Crick Institute receives its core funding from Cancer Research UK (FC001115), the UK Medical Research Council (FC001115) and the Wellcome Trust (FC001115). We would like to acknowledge Drs E. Grönroos, R. Hynds, S Ali and H Wood & Prof. P. Parker for their critical feedback on the manuscript.

## Author Contributions

W.G. Conceived the study, designed and carried out experiments, analyzed data and wrote the manuscript. A.R-M. Analyzed patient data. P.C-I. Carried out mass spectrometry analyses. J.J.M. Designed and carried out experiments. F.C. Analyzed patient data. A.P. Designed and carried out experiments. C.A.H. Supervised drug screening. P.C. Supervised mass spectrometry analyses. J.G. Provided patient samples and data. J.F. Provided patient samples and data. D.B. Conceived, write manuscript and supervised the study. All authors provided critical feedback on the manuscript pre-submission.

