## Supplementary Data for "CKS1-dependent proteostatic regulation has dual roles combating acute myeloid leukemia whilst protecting normal hematopoiesis"

**Supplementary Information**

**Supplementary Figures**

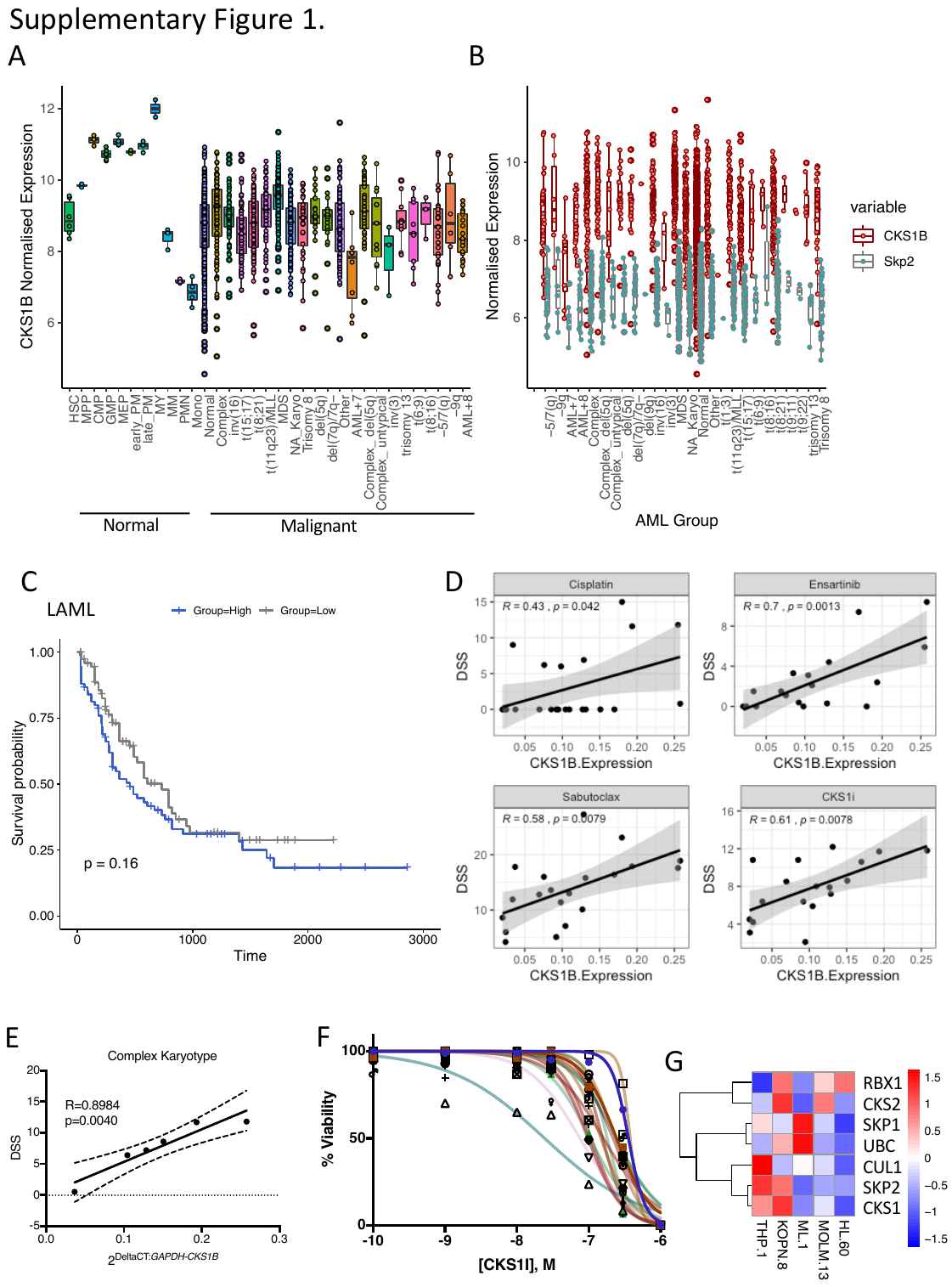

**Supplementary Figure 1. A.** *CKS1B* normalized expression and **B.** *SKP2* compared to *CKS1B* normalized expression of normal and malignant hematopoeitic cells obtained from Bloodspot.eu. Data sources: Human normal hematopoiesis (GSE42519), Human AML (GSE13159, GSE15434, GSE61804, GSE14468 and The Cancer Genome Atlas; TCGA). **C.** Overall survival of TCGA AML (LAML) patients stratified for *CKS1B* expression (50^th^ percentile). **D.** Correlation graphs between *CKS1B* expression and drug sensitivity for the indicated inhibitors in the poor risk AML cohort. 95% confidence interval and Pearson’s correlation coefficient for each drug are indicated. **E.** Correlation between complex karyotype patient AML CKS1i drug sensitivity (DSS) and *CKS1B* expression. 95% confidence intervals presented. Pearson’s correlation coefficient was calculated for correlation (R^2^) and significance (P; N=6). **F.** Percentage viability of all poor risk AML samples tested against a range of CKS1i concentrations (N=21). **G.** Expression of key SCF^SKP2-CKS1^ subunits in leukemic cell lines used in this study. Data presented are z-normalised (per gene) transcripts per million reads (TPMs) from the EBI Cell Line Expression Atlas.

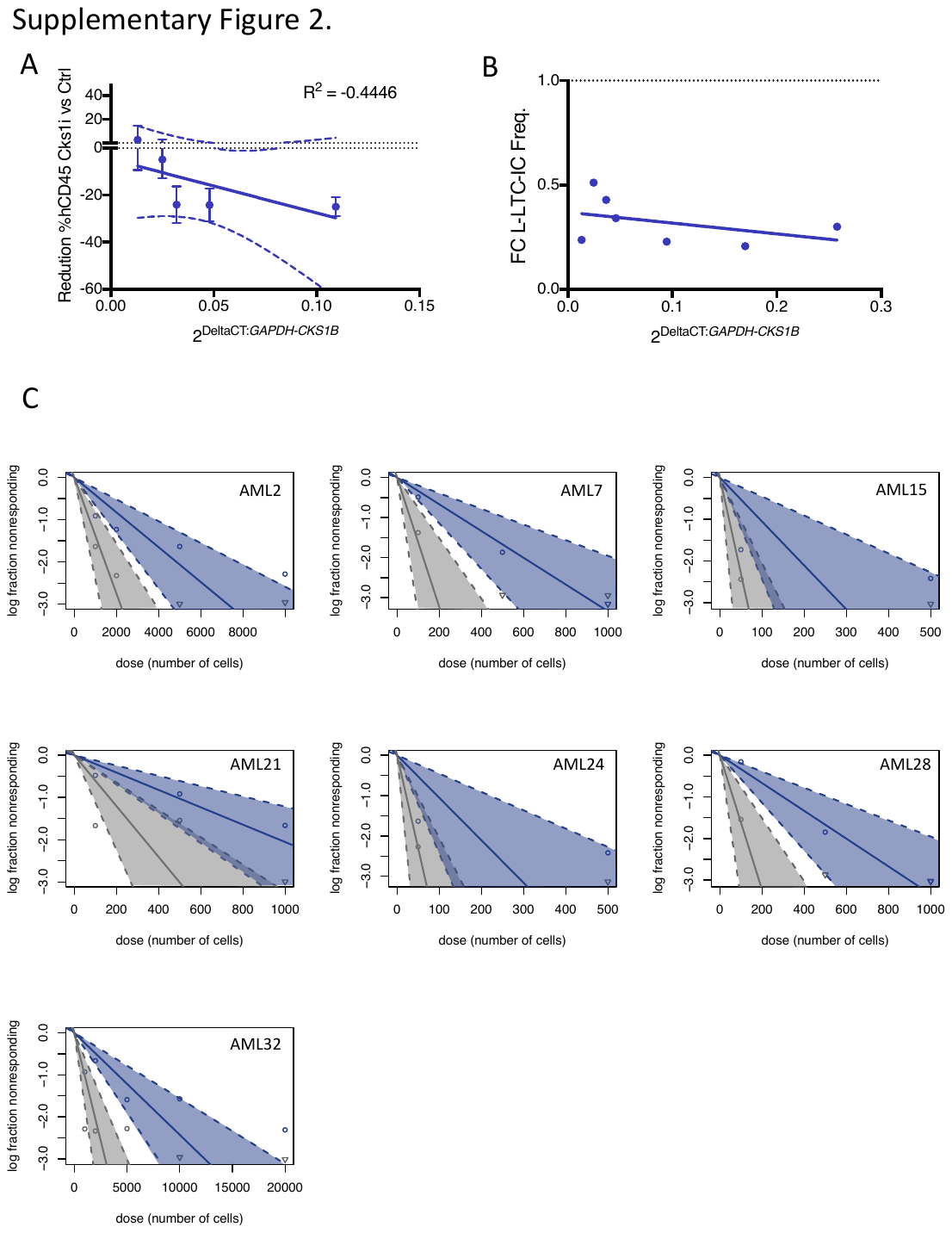

**Supplementary Figure 2. A.** Reduction in % AML burden in PDX, between CKS1i treatment versus control one-week post CKS1i treatment versus *CKS1B* expression. Pearson’s correlation coefficient was calculated for correlation (R^2^), 95% confidence interval shown. **B.** Fold change estimated L-LTC-IC frequency CKS1i treatment versus control at the end of the L-LTC-IC time period compared to *CKS1B* expression. **C**. Graph of estimated L-LTC-IC frequency for the indicated patients control and treated with CKS1i.

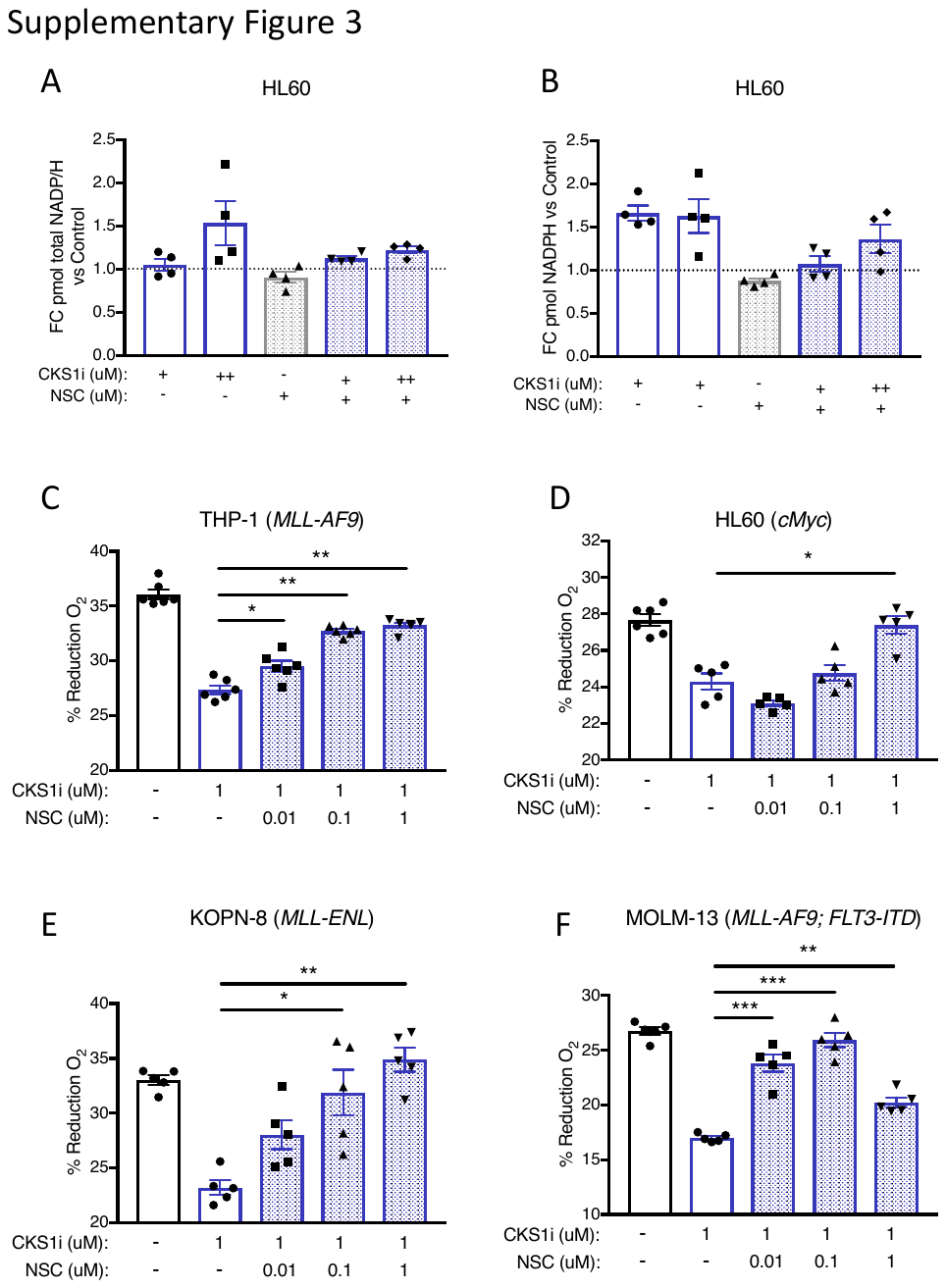

**Supplementary Figure 3. A.** Fold change pmol total NADP/NADPH in indicated treatments versus control (indicated dotted line at 1) for HL60 cells. CKS1i (+ = 1 μM, ++ = 5 μM) or NSC (+ = 0.1 μM) **B.** Fold change pmol NADPH in indicated treatments versus control for HL60 cells (N=4 per cell line and treatment). Viability represented by percentage reduction O_2_ of the indicated cell lines in response to the indicated concentrations of CKS1i and NSC23766 for **C.** THP-1, **D.** HL60, **E.** KOPN-8 and **F.** MOLM-13 cell lines (N=5 per cell line and treatment, except THP-1 where N=6).

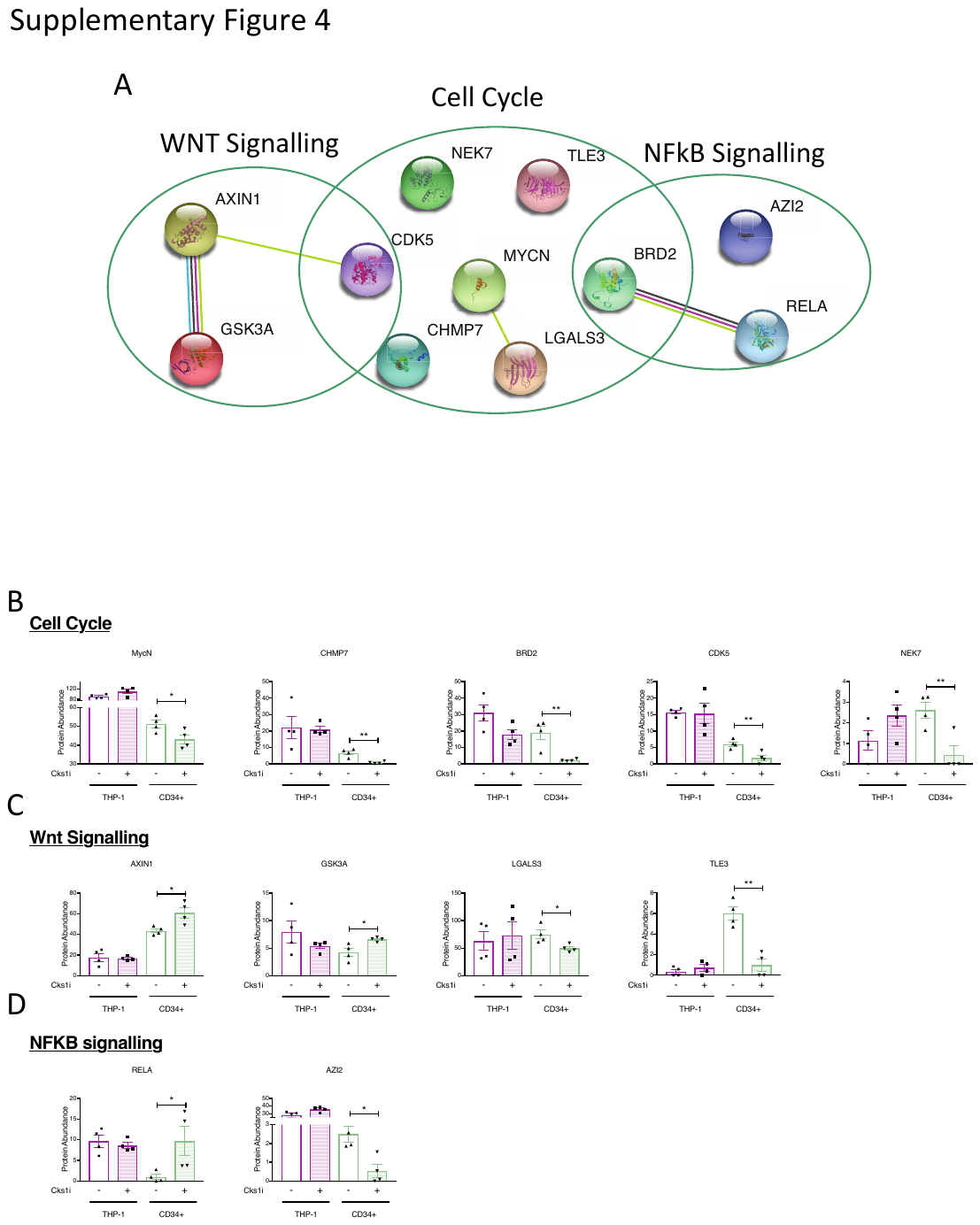

**Supplementary Figure 4. A.** String network connections for key pathways differentially regulated in CD34+ CKS1i treated cells. Protein abundances of **B.** cell cycle, **C.** Wnt signalling and **D.** NFkB signalling proteins differentially expressed in CD34^+^ CKS1i treated cells only (not THP-1 cells). * p<0.05; **p<0.01.

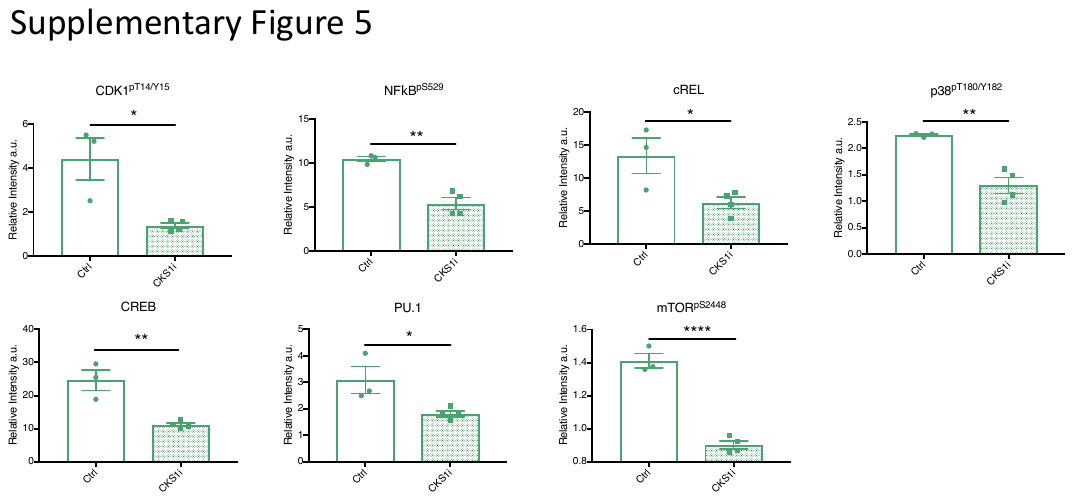

**Supplementary Figure 5.** Relative intensity of key protein markers significantly different between control and CKS1i treated CD34^+^ cells analysed by CyTOF. A Student’s *t*-test was used to calculate significance of difference for all graphs (Ctrl N=3, CKS1 N=4). * p<0.05; **p<0.01; ***p<0.001; ****p<0.0001.

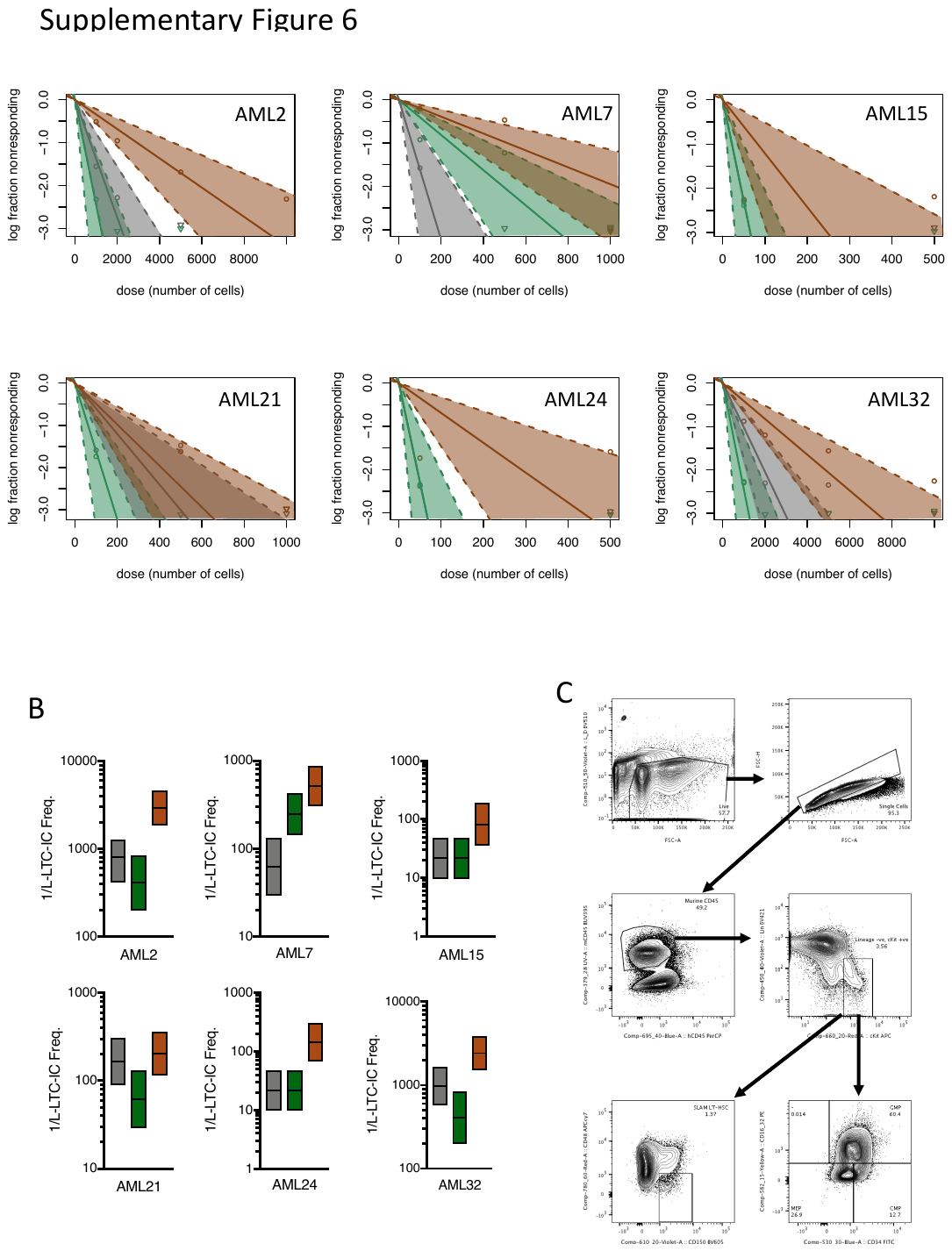

**Supplementary Figure 6. A.** Graph of estimated L-LTC-IC frequency for the indicated patients control (Grey) and treated with DA (Green) or DAC (Brown). **B.** Calculated L-LTC-IC frequencies and confidence intervals by ELDA (Control = Grey, DA = Green, DAC = Brown). **C.** Example gating strategy for residual murine LT-HSCs in xenograft models.

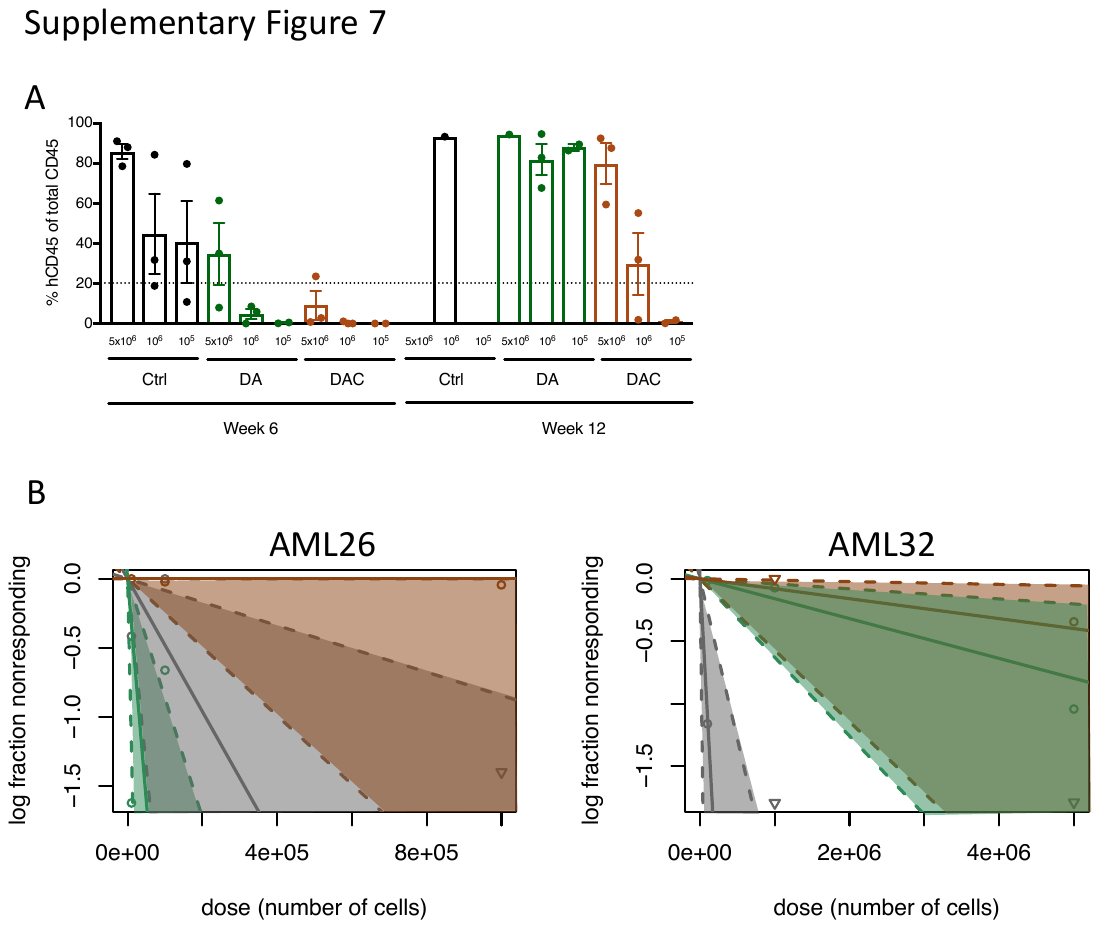

**Supplementary Figure 7. A.** Percentage hCD45 bone marrow engraftment of AML32 engrafted in secondary mice at limiting dilution weeks 6 and 12 (Ctrl N=3 per dose, DA 5x10^6^ & 10^6^ N=3 10^5^ N=2, DAC 5x10^6^ & 10^6^ N=3 10^5^). **B.** Graph of estimated LSC frequency for the indicated patients control (Grey) and treated with DA (Green) or DAC (Brown).

**Resources Table**

| **REAGENT or RESOURCE** | **SOURCE** | **IDENTIFIER** |
| --- | --- | --- |
| **Antibodies (FACS)** | | |
| hCD3 | BD | HIT3a |
| hCD19 | eBioscience | HIB19 |
| hCD33 | BD | WM53 |
| hCD34 | BD | 8G12 |
| hCD38 | eBioscience | HB7 |
| hCD45 | Biolegend | HI30 |
| hCD45RA | eBioscience | HI100 |
| hCD49f | BD | GoH3 |
| hCD90 | BD | _­­­_5E10 |
| Anti-Human Lineage Cocktail | BD | 340546 |
| mCD45 | eBioscience | 30-F11 |
| Anti-Mouse Lineage Cocktail | Biolegend | 133311 |
| mCD117 | BD | 2B8 |
| mSca1 | BD | D7 |
| mCD34 | BD | RAM34 |
| mCD48 | Biolegend | HM48-1 |
| mCD150 | Biolegend | TC15-12F12.2 |
| mCD16/32 | Biolegend | 93 |
| Annexin V | eBioscience | A-35122 |
| Ki67 | BD | 51-36524X |
| **Antibodies (CyTOF)** | | |
| CD45 (89Y) | Fluidigm | HI30 |
| CD33 (41Pr) | eBioscience | WM53 |
| P53pS392 (142Nd) | CST | 9281 |
| CD117 (143Nd) | Fluidigm | 104D2 |
| RBpS807/S811 (144Nd) | BD | J112-906 |
| SRCpY418 (145Nd) | BD | K98-37 |
| BCL2 (146Nd) | Fluidigm | EPR17509 |
| ELK (147Sm) | Abcam | ab28818 |
| B-Catenin (148Nd) | CST | D13A1 |
| PU.1 (149Sm) | CST | 9G7 |
| STAT5pY694 (150Nd) | Fluidigm | 47 |
| cRel (151Eu) | Abcam | ab30624 |
| AKTpS473 (152Sm) | Fluidigm | D9E |
| STAT1pY7001 (153Eu) | Fluidigm | 58D6 |
| P90-RSKpT359 (154Sm) | CST | D1E9 |
| CD34 (155Gd) | eBioscience | 4HI1 |
| P38PT180/Y182 (156Gd) | Fluidigm | D3F9 |
| STAT3pY705 (158Gd) | Fluidigm | 4/P-Stat3 |
| MAPKAPK2pT334 (159Tb) | Fluidigm | 27B7 |
| CD49f (160Gd) | eBiosciences | eBioGoH3 |
| BADpS112 (161Dy) | Fluidigm | 40A9 |
| CDK1pT14/Y15 (162Dy) | Life Technologies | 44-686G |
| GATA1 (163Dy) | CST | D52H6 |
| IkBa (164Dy) | Fluidigm | L35A5 |
| CREBpS133 (165Ho) | Fluidigm | 87G3 |
| NFkBpS529 (166Er) | Fluidigm | K10895.12.50 |
| ERK1/2pT202/Y204 (167Er) | Fluidigm | D13.14.4E |
| Ki67 (168Er) | Fluidigm | 3168001B |
| CD45RA (169Tm) | Fluidigm | HI100 |
| RET (170Er) | R&D systems | MAB718 |
| ZAP70pY319/SYKpY352 (171Yb) | Fluidigm | 17a |
| RETpY905 (172Yb) | CST | 3221 |
| CD90 (173Yb) | Fluidigm | 5E10 |
| mTORpS2448 (174Yb) | Life Technologies | 44-1125G |
| S6pS235/S236 (175Lu) | Fluidigm | N7-548 |
| CD38 (176Yb) | eBioscioence | HIT2 |
| **Chemicals, Peptides, and Recombinant Proteins** |  |  |
| hSCF | Peprotech | 300-07 |
| hFLT3L | Peprotech | 300-19 |
| hTPO | Peprotech | 300-18 |
| Saponin | Riedel-de Haen | 16109 |
| Cisplatin | Fluidigm | 201064 |
| CellRox Deep Red | Life Technologies | C10422 |
| Click-IT OP-Puromycin Kit | Life Technologies | C10456 |
| NSC23766 | Selleckchem | S8031 |
| SKP2E3LI (a.k.a. CKS1i) | Merck | 500519 |
| **Mouse models (inc. *in vivo* N numbers)** |  |  |
| NOD-SCID IL2Ry^-/-^ (NSG) | The Jackson Laboratory | N/A |
| Figure 1. THP-1 | NSG | 6 Ctrl, 7 CKS1i |
| Figure 1. HL60 | NSG | 5 Ctrl, 6 CKS1i |
| Figure 2. AML12 | NSG | 7 Ctrl, 7 CKS1i |
| Figure 2. AML21 | NSG | 5 Ctrl, 6 CKS1i |
| Figure 2. AML26 | NSG | 7 Ctrl, 7 CKS1i |
| Figure 2. AML27 | NSG | 6 Ctrl, 7 CKS1i |
| Figure 2. AML32 | NSG | 6 Ctrl, 6 CKS1i |
| Figure 4. CD34+ | NSG | 5 Ctrl, 3 DA, 3 DAC |
| Figure 5. AML12 | NSG | 6 Ctrl, 7 DA, 7 DAC |
| Figure 5. AML21 | NSG | 10 Ctrl, 10 DA, 11 DAC |
| Figure 5. AML26 | NSG | 7 Ctrl, 7 DA, 7 DAC |
| Figure 5. AML32 | NSG | 10 Ctrl, 9 DA, 9 DAC |
| Figure 5. AML26 Secondary Transplantation | NSG | 7 Ctrl, 7 DA, 4 DAC |
| Figure 5. AML32 Secondary Transplantation | NSG | 9 Ctrl, 8 DA, 8 DAC |
| **Oligonucleotides** |  |  |
| Oligo dT_20_ | Sigma Aldrich | N/A |
| Gggaaggtgaaggtcggagt | Sigma Aldrich | GAPDH_F |
| Gggtcattgatggcaacaata | Sigma Aldrich | GAPDH_R |
| CACCATTGGCAATGAGCGGTTC | Sigma Aldrich | B-ACTIN_F |
| AGGTCTTTGCGGATGTCCACGT | Sigma Aldrich | B-ACTIN_R |
| ATGTCTGAATCTGAATGGAGG | Sigma Aldrich | CKS1_F |
| TCATTTCTTTGGTTTCTTGGG | Sigma Aldrich | CKS1_R |
| **Software, Algorithms and Data availability** |  |  |
| R software v3.6.1 | R project | r-project.org |
| CytoBank | MRC | mrc.cytobank.org |
| FlowJo v10.6.1 | FlowJo | N/A |
| Prism 7 | GraphPad | N/A |
| Figure 3 proteomic analyses | PRIDE |  |
| **Primary samples and cell lines** |  |  |
| Patient AML samples | Bart’s Biobank | N/A |
| Human Umbilical Cord Blood | Royal London Hospital | N/A |
| THP-1 | The Francis Crick Cell Services | N/A |
| HL60 | The Francis Crick Cell Services | N/A |
| ML-1 | The Francis Crick Cell Services | N/A |
| MOLM-13 | The Francis Crick Cell Services | N/A |
| KOPN-8 | The Francis Crick Cell Services | N/A |
